# Domoic acid biosynthesis and genome expansion in *Nitzschia navis-varingica*

**DOI:** 10.1101/2025.04.19.649686

**Authors:** Steffaney M. Wood-Rocca, Nicholas Allsing, Yasuhiro Ashida, Masaki Mochizuki, Malia L. Moore, Zoltán Füssy, Yuichi Kotaki, Clyde Puilingi, Yukari Maeno, Aodhan W. Beattie, Andrew E. Allen, Mari Yotsu-Yamashita, Todd P. Michael, Bradley S. Moore

**Affiliations:** Center for Marine Biotechnology and Biomedicine, Scripps Institution of Oceanography, University of California San Diego, La Jolla, CA; Environmental Genomics group, J. Craig Venter Institute, La Jolla, CA; The Plant Molecular and Cellular Biology Laboratory, Salk Institute for Biological Sciences, La Jolla, CA; Graduate School of Agricultural Science, Tohoku University, Aramaki-Aza-Aoba, Aoba-ku, Sendai, Japan; Faculty of Science and Technology, Solomon Islands National University, Honiara, Solomon Islands; School of Science & Technology, Pacific Adventist University, Private Mail Bag, Boroko, NCD, Papua New Guinea; Graduate School of Agricultural and Life Sciences, The University of Tokyo, Yayoi, Bunkyo-ku, Tokyo, Japan; Integrative Oceanography, Scripps Institution of Oceanography, University of California San Diego, La Jolla, CA; Skaggs School of Pharmacy and Pharmaceutical Sciences, University of California San Diego, La Jolla, CA

## Abstract

Production of the neurotoxin domoic acid (DA) by benthic diatom *Nitzschia navis-varingica* poses considerable health and economic concerns. In this study, we employed whole genome sequencing and transcriptomic analyses of regionally distinct *N. navis-varingica* strains to unravel the genomic underpinnings of DA biosynthesis. Our analyses revealed sizable genomes—characterized by an abundance of repetitive elements and noncoding DNA—that exceed the size of any other pennate diatoms. Central to our findings is the discovery of an expanded domoic acid biosynthesis (*dab*) gene cluster, spanning over 60 kb and marked by a unique organization that includes core genes interspersed with additional genetic elements. Phylogenetic and syntenic comparisons indicate that transposition events may have driven the expansion and reorganization of this cluster. Biochemical assays validated that the kainoid synthase encoded by *dabC* catalyzes the formation of isodomoic acid B, thereby establishing a distinct chemotype in contrast to the DA profiles of planktonic diatoms. These results highlight the evolutionary trajectory of DA biosynthesis in diatoms and potential advantages conferred by genome expansion and enzyme diversification in dynamic marine environments.

**IMPORTANCE:** Domoic acid (DA) is a potent neurotoxin produced by marine micro- and macroalgae problematic to fisheries and toxic to humans and animals. Our study elucidates the molecular mechanisms underlying DA production in the widespread Western Pacific benthic diatom, *Nitzschia navis-varingica*. Genomic and biochemical insights add information to our understanding of the evolution of toxin production across diverse phyla and also fill a gap in the knowledge of secondary metabolism in marine diatoms. These findings provide a genetic framework for identifying toxin production and its impacts in the benthos of vulnerable, coastal ecosystems.

## INTRODUCTION

Domoic acid (DA) is a member of the kainoid class of natural neurotoxins, which includes the isomers and derivatives of DA, the namesake kainic acid, and acromelic acid (1, 2). Kainoids are non-proteinogenic amino acids characterized by a glutamate-derived pyrrolidine ring and multiple carboxylates (Fig. 1A). DA functions as a potent glutamate receptor agonist, leading to neurotoxic effects such as memory loss, disorientation, vomiting, seizures, and in severe cases, coma and death (3–5). The first recorded outbreak of DA poisoning in humans, termed amnesic shellfish poisoning, occurred in 1987 in Prince Edward Island, Canada. This outbreak was attributed to the accumulation of DA in mussels that had consumed the toxin-producing diatom from what is now known as *Pseudo-nitzschia multiseries* (6, 7).

**Figure 1.**
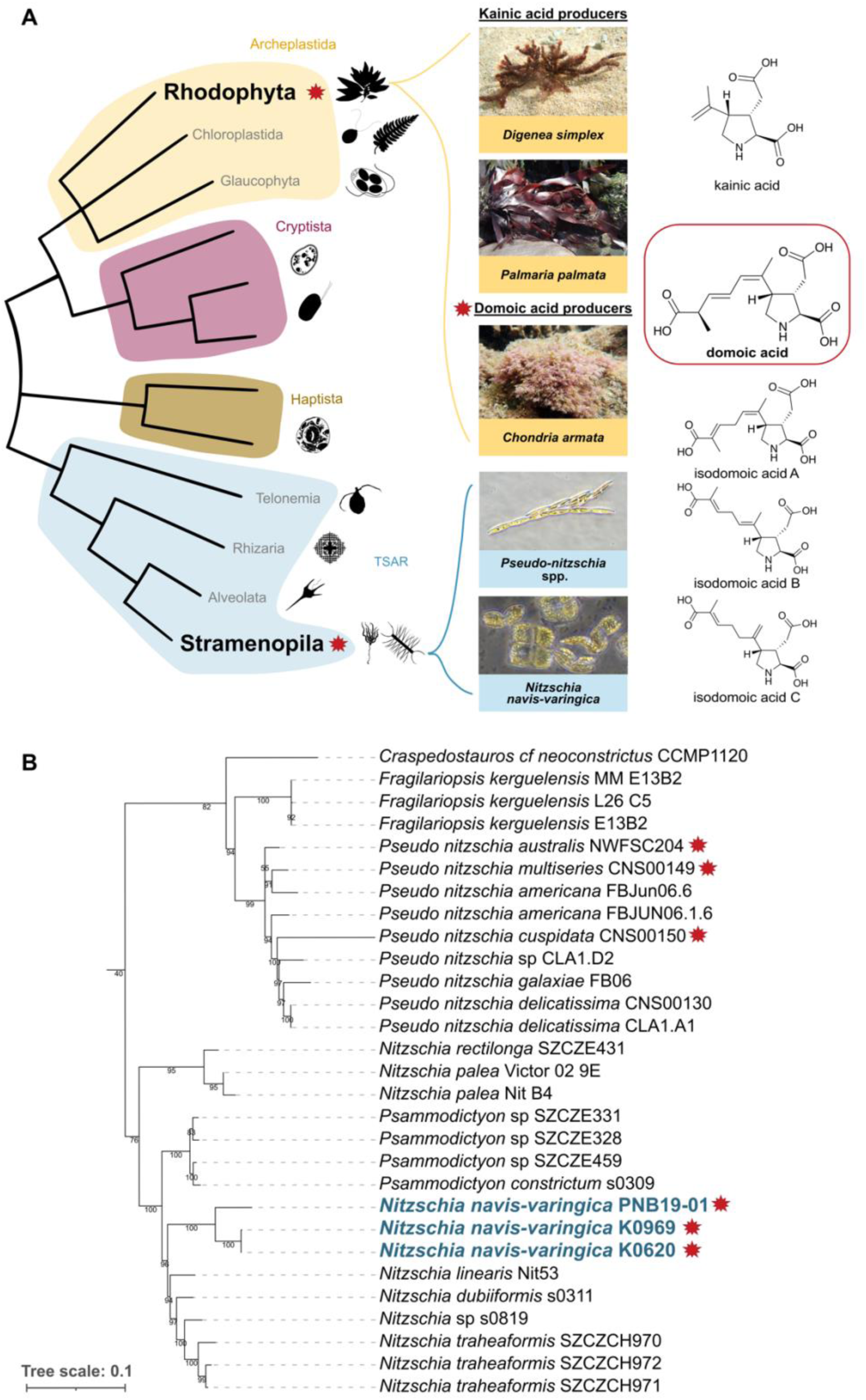
Kainoid production across marine eukaryotes. (A) Simplified eukaryotic tree of life based on Burki et al. with DA and kainic acid producers highlighted within their respective supergroups (38). (B) Rooted multi-locus phylogenetic analysis of alignments of 18S (SSU) and 23S (LSU) nuclear RNA genes as well as chloroplast genes *psbC* and *rbcL* from the Bacillariaceae family. Supplementary Figure S1 shows a complete version of this phylogeny. Stars indicate *dab* cluster identified from this species. Bold indicates strains highlighted in this study.

Since this deadly initial outbreak, subsequent research has described DA production in a variety of marine eukaryotic algae (8–11). Approximately half of the over 50 described species of harmful algal bloom-forming diatoms in the genus *Pseudo-nitzschia* have been reported to produce DA (12). These toxic blooms have caused significant negative economic impact and safety risks to humans and animals through bioaccumulation of DA in the marine food web (12–14). Additionally, red macroalgae of the family Rhodomelaceae—most notably *Chondria armata*, from which DA was first described—and other pennate diatoms such as *Amphora* sp., *Nitzschia bizertensis*, and *Nitzschia navis-varingica* have also been demonstrated to produce this neurotoxin (9, 12, 15).

*N. navis-varingica* is commonly found in benthic ecosystems in the Western Pacific, where its DA production poses a threat to aquaculture facilities and shellfish harvested from mangrove ecosystems (16, 17). DA production in *N. navis-varingica* is widespread, corresponding with its ability to inhabit diverse ecological niches in conditions ranging from planktonic to benthic and euhaline to brackish systems (18–20). *N. navis-varingica* strains tend to produce higher ratios of isodomoic acid B to DA, in contrast to *Pseudo-nitzschia* spp. that primarily produce DA and isodomoic acid A (20, 21). Recent studies have demonstrated that geographically distinct populations display a high degree of genetic homogeneity and strains with similar DA profiles are typically monophyletic, underscoring a genetic determinant in toxin production (17, 19, 22).

With the expansion of whole genomic sequencing of DA-producing algae, the evolution of DA biosynthesis is being unraveled (15, 23). DA biosynthesis genes (*dab*) were first identified in *P. multiseries* in a biosynthesis gene cluster (BGC) encoding a *N*-prenyltransferase (*dabA*), hypothetical protein (*dabB*), kainoid synthase (*dabC*), and cytochrome P450 (CYP450, *dabD*) (23). *Dab* gene clusters have thus far been identified in toxigenic *Pseudo-nitzschia* spp., including *P. multiseries*, *P. australis*, *P. seriata*, *P. cuspidata*, and *P. multistriata*, with their protein sequences displaying high amino acid identity (23–27).

DA and kainic acid biosynthesis have also been described in red macroalgae *Chondria armata* as well as *Digenia simplex* and *Palmaria palmata*, respectively (15, 28, 29). Despite the evolutionary distance between diatoms (Bacillariophyta) and red algae (Rhodophyta), the red algal domoic acid biosynthesis genes (*radACD*) display a high degree of gene synteny with both kainic acid biosynthesis genes (*kabAC*) and the DA biosynthesis (*dabABCD)* gene clusters of *Pseudo-nitzschia* species (15). The unique structure and activity of these enzymes has led to additional research, as they represent a possible horizontal gene transfer event across distant taxa and novel biochemical mechanisms and enzyme structures (30–33). These discoveries highlight the complex evolutionary history of DA biosynthesis and underscore the need for further research.

Despite the incredible diversity within *Nitzschia*, comprising over 170,000 species and intraspecific names, only a handful of non-DA producers have been sequenced (34–36). Genomic studies of benthic diatoms such as *N. inconspicua* and *Seminavis robusta* revealed expansion of repetitive elements and unique adaptations to the benthos (35, 37). We therefore suspect *N. navis-varingica* will show a similar structure in line with its benthic adaptations. Moreover, these genomic data are essential for characterizing the evolution of the *dab* pathway in a novel lineage of algae.

To further investigate the evolution of DA biosynthesis and the ecological adaptations of sub/tropical benthic diatoms, herein we sequenced regionally distinct strains of *N. navis-varingica* for comparative genome and transcriptome analyses and DA biochemical validation. We report here that genome expansion in *N. navis-varingica* is mirrored in the enlarged organization of the *dab* BGC that encodes the synthesis of the intermediate isodomoic acid B, in contrast to *Pseudo-nitzschia* diatoms, thereby suggesting a distinct evolutionary history in these diatom genera.

## RESULTS

### *Nitzschia navis-varingica* molecular taxonomy and genome characteristics

*N. navis-varingica* strains K0620 and K0969 were isolated from a marine shrimp culture pond and coastal waters in northern Vietnam, respectively (39). Strain PNB19-01 originated from Bootless Bay near Port Moresby, Papua New Guinea, following previous reports of DA and isodomoic acid B production (19). We investigated their molecular taxonomy within the Bacillariaceae family using a multi-locus phylogeny (18S rRNA, 23S rRNA, *psbC*, and *rbcL*). All three strains formed a well-supported, monophyletic lineage within a clade containing *N. linearis*, *N. dubiformis*, and *N. traheaformis*, remaining distinct from the DA-producing *Pseudo-nitzschia* clade (Fig. 1B). The *N. navis-varingica* clade is most closely related to *Psammodictyon* species, indicating that *N. navis-varingica* belongs to a separate lineage outside the major *Nitzschia* clades. The *Nitzschia* genus in this phylogeny is polyphyletic (Fig. S1), as has been previously reported and further evidenced by the clustering of *Nitzschia* spp. and *Psammodictyon*. Indeed, Mann et al. (2021) suggest that the two clades harboring *N. navis-varingica* may merit recognition as new genera, given the lack of clear morphological synapomorphies to unite them under *Nitzschia* as currently defined (34).

We produced partially phased genome assemblies of K0620 and K0969 and transcripts for PNB19-01 to investigate diversity in *dab* genes across DA-producing organisms as well as regionally distinct *N. navis-varingica* isolates. For K0620 and K0969, we used PacBio genomic and IsoSeq transcriptomic sequencing, while for PNB19-01, we used Novogene RNA-seq. PacBio sequencing produced HiFi reads to ∼35× coverage of K0620 with a read N50 of 16,137 bp, and ∼31× coverage of K0969 with a read N50 of 13,179 bp. IsoSeq transcriptomic sequencing resulted in 13,042,775 reads with an N50 of 1,415 bp for K0620 and 11,434,165 reads with an N50 of 1,537 bp for K0969. Analysis of K-mer composition of the HiFi reads reveals distinct homozygous and heterozygous peaks as well as elevated duplication, highlighting the diploid nature of the genome and the pronounced lower coverage heterozygous peak (Fig. S2). Assembling K0620 and K0969 HiFi reads into haploid genomes produced Haplotype 1 and Haplotype 2 assemblies for each, and while both assemblies are available, we use the Haplotype 1 for our analyses due to higher quality (Table S1). Haplotype 1 of K0620 assembled to 879.5 Mbp with an N50 of 303,884 bp and K0969 to 782.8 Mbp with an N50 of 134,724 bp (Table 1). For PNB19-01, Novogene RNA-seq resulted in 12,640,360 reads, 3.79 Gbp, with an N50 of 1,759 bp.

**Table 1.**
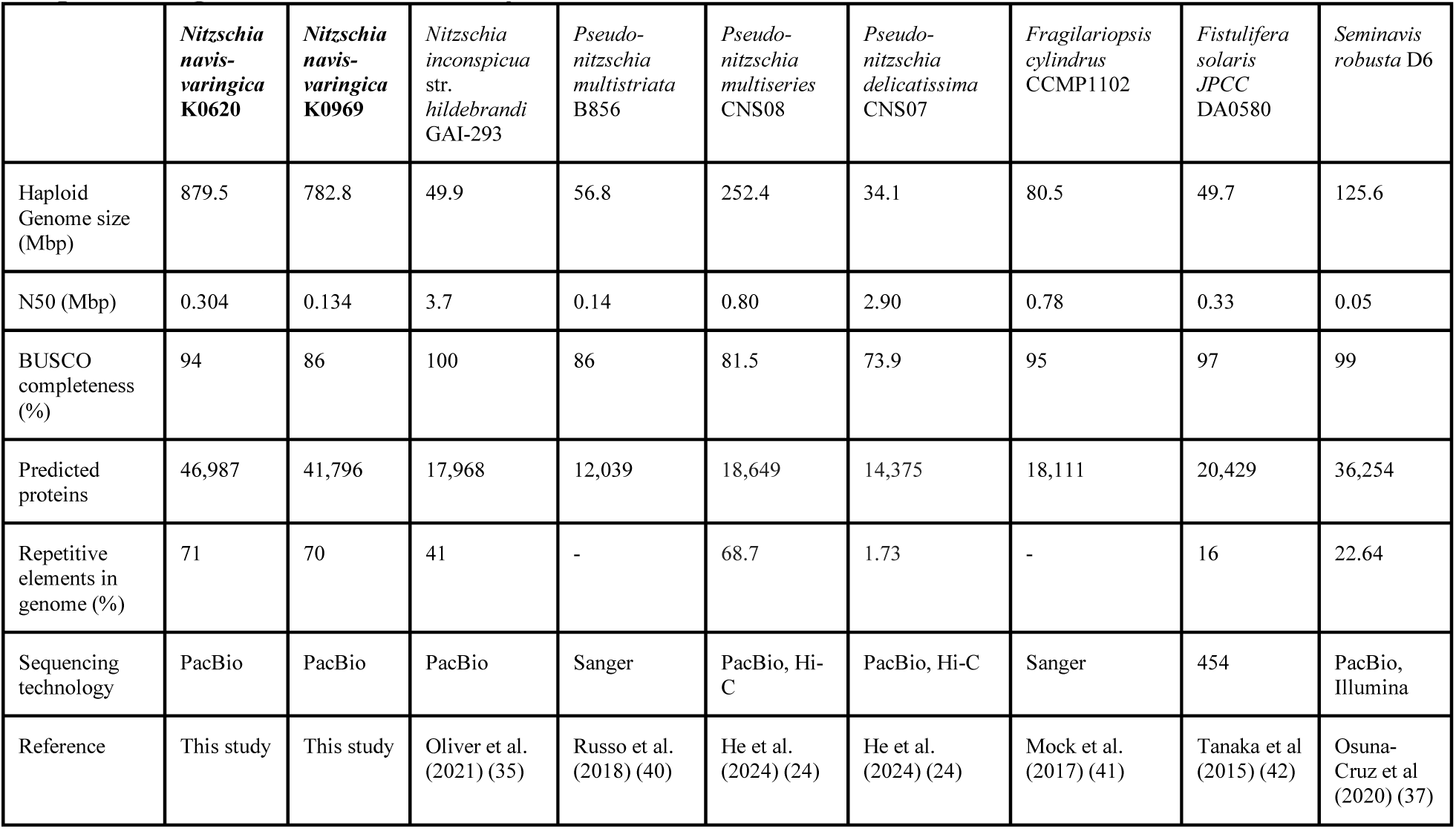
Genome characteristics of select pennate diatoms. See Table S1 for complete statistics for genomes produced in this study.

Both genomes are at least 15 times the size of *Nitzschia inconspicua* str. *hildebrandi* GAI-293 and up to 25 times the size of other pennate diatoms such as *P. delicatissima*. Despite the difficulty removing bacterial contamination prior to sequencing, the quality of assemblies was high, as evidenced by conserved Stramenopiles marker genes with 95% and 85% BUSCO completeness for K0620 and K0969, respectively (Fig. 2A, Table 1). Comparison of GC content with sequencing depth supports minimal contamination given that the majority of contigs cluster around the average GC content of approximately 33% (Fig. S3). Indicative of allelic variation retained in the assemblies, BUSCO analysis also identified a duplication of 19% and 31% for K0620 and K0969, respectively. Moreover, IsoSeq mapping statistics indicated that 91.5% and 83% of transcriptomic reads aligned to the K0620 and K0969 assemblies, further supporting the completeness of the gene models.

**Figure 2.**
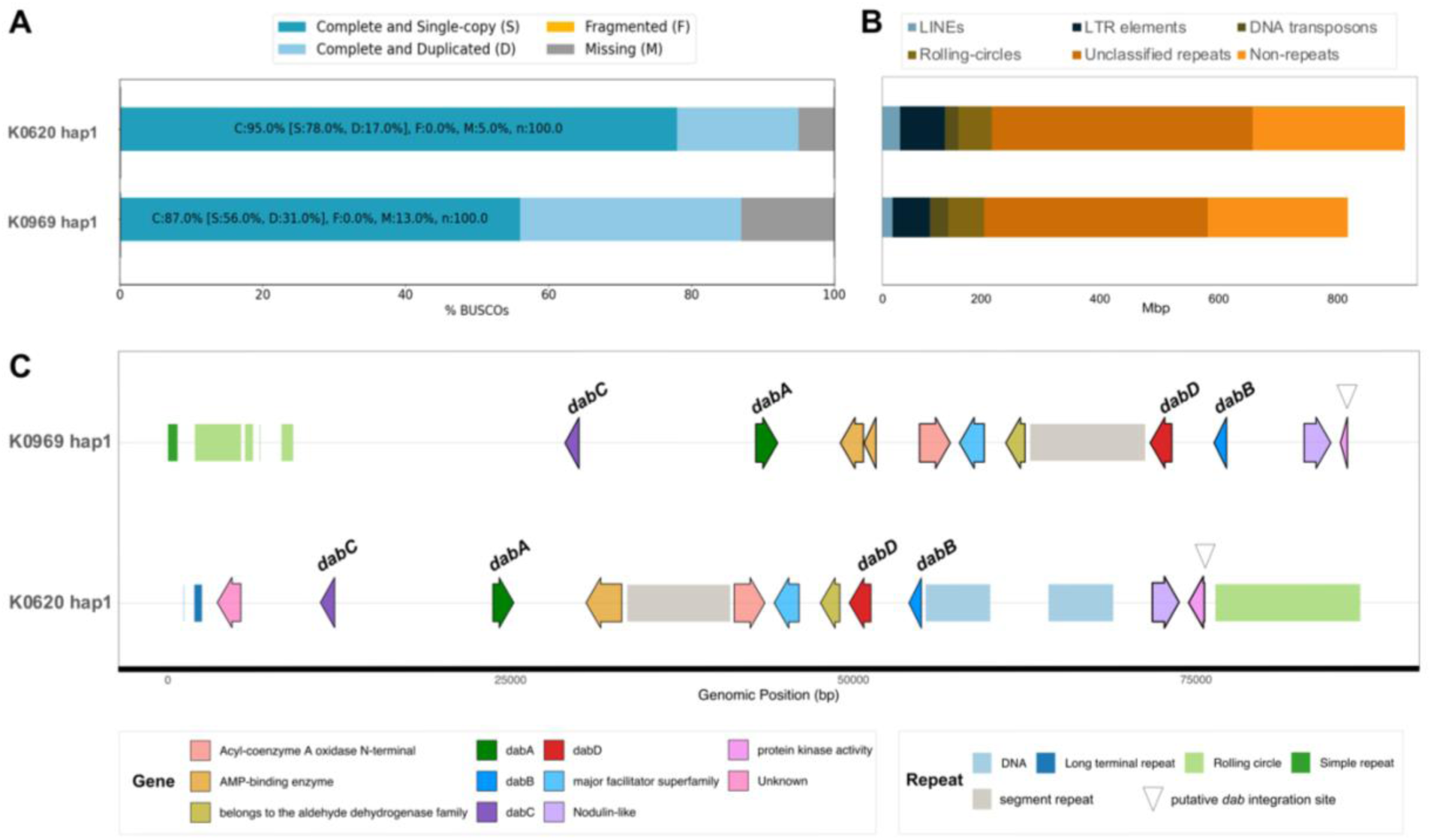
Genome Assembly Quality and Repeat Composition Across Haplotypes. **(A)** BUSCO completeness assessments highlighting the proportion of complete single-copy (S), duplicated (D), fragmented (F), and missing (M) orthologs in each assembly. **(B)** Length of different repeat elements within genome assemblies of K0969 and K0620. Categories include long interspersed nuclear elements (LINEs), long terminal repeat (LTR) elements, DNA transposons, rolling-circles, unclassified repeats, and non-repeats. See haplotype genome and repeat statistics in Table S1. **(C)** Genomic neighborhood of the *dab* gene cluster. Arrows represent genes and bars represent repetitive elements. See full length *dab*-containing contigs in Fig. S3.

The genome size in *N. navis-varingica* is shaped by the large proportion of non-protein coding regions and repetitive DNA (Fig 2B). Both strains only constitute approximately 7% coding sequences, in contrast to approximately 70% repetitive DNA. Despite this, they still contain more predicted proteins than other pennate diatom genomes (Table 1). Of the various classes of repetitive DNA, unclassified repetitive elements make up the largest fraction of the genomes (48-50%), followed by long-terminal repeats (LTR) (8-9%) and rolling-circles (6-8%). A similar makeup has been previously reported in centric diatom genomes (43).

Of the functionally annotated genes, the majority correspond to core metabolic processes (Fig. S4). The prevalence of annotations related to macromolecule transport, such as bicarbonate, nitrate, and magnesium, light harvesting, and stress response possibly reflect adaptations to the environmental fluctuation of pH, light and nutrient availability, and temperature that is common in the sub/tropical benthos (44–46). Given the ability of benthic *Nitzschia* spp. to produce volatile, halogenated hydrocarbons, like methyl iodide (47–49), we queried the genomes for putative haloperoxidase and halide methyltransferase genes. We observed an interesting colocalization of genes encoding a putative halide methyltransferase with an iodotyrosine deiodinase, the latter of which had homologs in benthic diatom *Seminavis robusta* and various Alveolates (Table S2, Fig. S5). The *N. navis-varingica* iodotyrosine deiodinase may function to detoxify halogenated amino acids, which in turn would result in cellular iodide that would be methylated and removed as volatile methyl iodide (49–53).

### Domoic acid biosynthesis gene cluster identification

We identified the *dab* gene cluster in the genomes of *N. navis-varingica* K0620 and K0969 using BLAST and HMM searches with the existing collection of *Pseudo-nitzschia dab* genes and *C. armata* red algal domoic acid (*rad*) genes. Initial searches revealed the co-localization of homologs for *dabA*, *dabB*, *dabC*, and a CYP450 unrelated to canonical *dabD* genes in both K0620 and K0969 (Fig. 2 and S5). An additional copy of *dabC* was also identified in K0620 (referred to as *dabC2*) on a separate contig (Fig S7). Notably, we identified a similar set of *dabABC* and a CYP450 transcripts in *N. navis-varingica* PNB19-01. Curiously, the CYP450 sequences that displayed the strongest homology with *Pseudo-nitzschia dabD* sequences were not co-localized with *dabABC* in K0620 and K0969. Rather, the co-localized CYP450s only exhibited up to 21% amino acid identity with known *dabD* sequences, a trend also observed in *C. armata* (15, Fig. S8). We thus suspect that the co-clustered CYP450, which is conserved across all three strains, functions as the oxygenase DabD. Additionally, the *N. navis-varingica dab* cluster contains the *dabB* gene that encodes a protein of unknown function as first reported in *Pseudo-nitzschia* diatoms, yet not in DA-producing red algae (23).

The *N. navis-varingica dab* cluster spans over 60 kb, features a unique gene arrangement, and contains additional genes and repetitive elements (Fig. 2C). The expansion of intergenic space and the abundance of repeats at the *dab* locus mirror the overall genome size and composition. The cluster is flanked by DNA transposons and rolling circle elements, suggesting that transposition events may have contributed to its expansion and rearrangement. Within the cluster, interspersed genes encode proteins with domains such as aldehyde dehydrogenase, major facilitator superfamily, acyl-coenzyme A oxidase, and AMP-binding enzymes. To further determine if these genes were potential segment repeat elements misannotated by RepeatMasker (v4.1.8), the intergenic space between *dabA* and *dabD* was BLAST searched against the remainder of the genome assembly (54). Results showed that portions of the non-coding intergenic space, but not the inserted coding sequences, displayed high (up to 95%) nucleotide similarity throughout the genome (Fig. S9).

The gene order of the *N. navis-varingica dab* cluster is rearranged relative to other DA-producing organisms: *dabA* and *dabC* are adjacent, followed by a series of interspersed genes, then *dabB* and the putative *dabD*. The CoA-binding protein and protein kinase genes that flank a block containing the aldehyde dehydrogenase, major facilitator superfamily member, *dabB*, *dabD*, and a nodulin-like gene are similar to the sequences found in other diatom genomes.

The putative integration site for the *Pseudo-nitzschia dab* gene cluster, involving a highly conserved CoA-binding protein (K0620 hap1 g25680) and protein kinase ( K0620 hap1 g25681) gene pair as proposed by He et al., is present in *N. navis-varingica* in a genomic region separate from the *dab* cluster (24). Interestingly, genes containing similar domains are found in the *N. navis-varingica dab* locus. The protein kinase gene found downstream of the *N. navis-varingica dab* cluster is more homologous (31% amino acid identity) to the sequence found in putative *Pseudo-nitzschia dab* insertion site than the CoA-binding gene (19% amino acid identity). Phylogenetic analysis of these sequences shows that these sequences are highly conserved across diatoms (Fig S10). This observation suggests that the putative integration site may be highly conserved across many diatom lineages. The presence of orthologous protein kinase may indicate a possible conserved mechanism of cluster integration across these taxa.

### *N. navis-varingica* compound detection and biochemical validation of isodomoic acid B synthase

Analysis of *N. navis-varingica* culture extracts using liquid chromatography mass spectrometry (LCMS) revealed the presence of DA along with related isomers isodomoic acid A and isodomoic acid B (Fig. 3 and S11). While *N. navis-varingica* cultures primarily produce DA, isodomoic acid B is present at substantial levels—a pattern that contrasts with DA-producing *Pseudo-nitzschia* species that predominantly yield DA and isodomoic acid A. Based on these observations, we hypothesized that the kainoid synthase enzyme (DabC) in *N. navis-varingica* generates isodomoic acid B, a pattern found in isodomoic acid B chemotype seen in *C. armata* (15).

**Figure 3.**
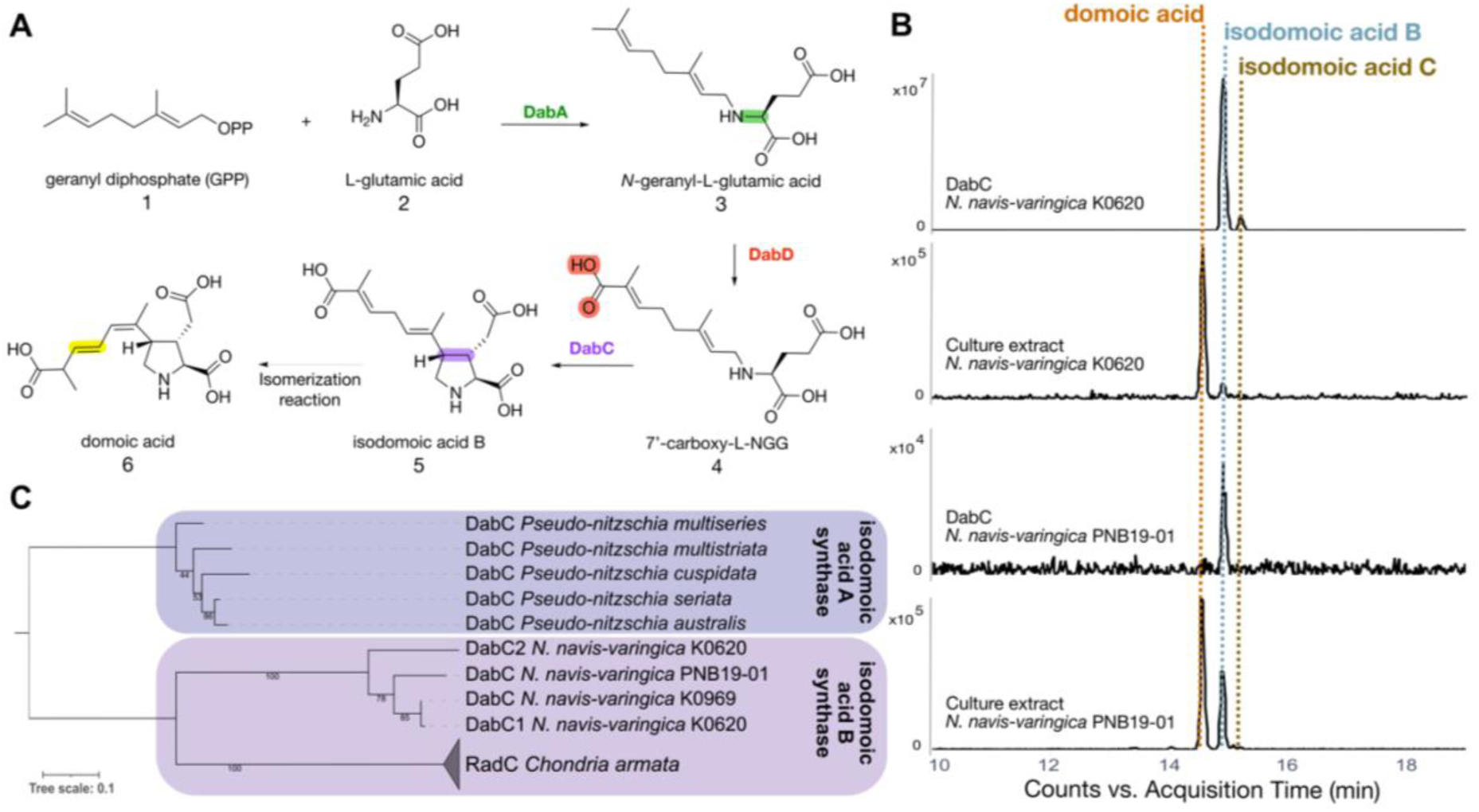
Domoic acid biosynthesis pathway in *N. navis-varingica* proceeds through isodomoic acid B. **(A)** *N. navis-varingica* Dab enzymes collectively L-glutamic acid (L-Glu) and geranyl diphosphate (GPP) to isodomoic acid B on route to DA. **(B)** Extracted ion chromatogram profiles for DA and isodomoic acid B (*m/z* 312.1±1.0) from DabC assays and culture extracts. See comparison to synthetic standards in Fig. S7. **(C)** Branching of isodomic acid B synthase DabC amino acid sequences visible through unrooted phylogeny of kainoid synthase enzymes. Clade shading indicates isodomoic acid A or B production.

To validate this hypothesis, we examined key steps in the DA biosynthesis pathway. Enzyme assays with purified glutamate *N*-prenyltransferase enzymes (K0620-NnvDabA and PNB19-01-NnvDabA) confirmed the production of *N*-geranyl-L-glutamic acid (Fig. 3, compound **3**, L-NGG) from geranyl diphosphate (**1**) and L-glutamate (**2**, Fig. S12). Subsequent assays of the kainoid synthase enzymes K0620-NnvDabC and PNB19-01-NnvDabC revealed that they catalyzed the oxidative cyclization of linear precursor 7’-carboxy-L-NGG (**4**), yielding isodomoic acid B (**5**) as the product (Fig. 3B). These results establish that *N. navis-varingica* exhibits a distinct chemotype from other DA-producing diatoms in its production of isodomoic acid B. It also underscores the role of the kainoid synthase in driving isomer diversity within the DA biosynthesis pathway.

NnvDabC enzymes also cyclize L-NGG (**3**) to produce primarily dainic acid A (Fig. S13). NnvDabC from strain K0620 also appears to produce trace amounts of dainic acid B/C, which have been shown to co-elute using our methods (15, 23). This is consistent with enzymatic products produced by isodomoic acid B synthase RadC and *P. multiseries* isodomoic acid A synthase DabC, but a marked discrepancy from KabC, which cannot cyclize the geranyl side chain present *dab* pathway intermediates (14). It should also be noted that dainic acid A can sometimes be detected in culture extracts of *N. navis-varingica* (Fig. S14), as has been previously reported in *C. armata* (29).

### Gene synteny and phylogenetic analysis of kainoid biosynthesis gene clusters

Phylogenetic analysis of individual *dab* genes largely aligns with previous studies (15, 23, 28, 30). Isodomoic acid B synthase NnvDabC groups more closely with RadC, rather than with the isodomoic acid A synthases found in more closely related *Pseudo-nitzschia* species (Fig. 3C). All kainoid synthase DabC enzymes nonetheless form a subclade in the broader phylogeny of bacterial and fungal alpha-ketoglutarate dependent Fe(II)-containing dioxygenases as previously observed (15), highlighting the sources of a likely horizontal gene transfer event (Fig. S15A). DabB phylogeny shows that NnvDabB groups separately from *Pseudo-nitzschia* spp. DabB sequences, and other uncharacterized proteins sequences in the tree are distantly related (Fig. S16). In silico analysis of NnvDabB indicates that it contains N-terminal signal peptide and may act as a type II signal anchor that anchors proteins to the membrane (55, 56)

Broader phylogeny of DabD sequences within CYP450s highlights the uniqueness of *N. navis-varingica* DabD that groups with other diatom CYP450 sequences but separately from *Pseudo-nitzschia* spp. DabD, also echoing the possibility of gene duplication and lineage-specific neofunctionalization of CYP450 enzymes as reported in the *rad* pathway (15, Fig. S15B). To understand the evolutionary origin of the taxonomically distinct NnvDabD co-localized with the *dab* cluster, we constructed a phylogenetic tree of all 128 annotated K0620 CYP450 sequences (Fig. S17). While the *dabD* gene candidate predicted by homology to the characterized *Pseudo-nitzschia* spp. *dabD* formed a clade with CYPs of mostly unknown function, the *dabD* candidate from the BGC groups with CYP450s widely annotated as retinoid hydroxylases. This result indicates a possible origin of the candidate CYP450 DabD in carotenoid biosynthesis, potentially resulting from similarities in structural motifs of the two isoprenoid species.

In the larger context of kainoid BGCs, a concatenated phylogeny of *N*-prenyltransferase (A-type), B proteins (unknown function), kainoid synthase (C-type), and CYP450 (D-type) genes forms a well-supported *N. navis-varingica* clade distinct from both the *Pseudo-nitzschia dab* clusters and the red algal (Rhodophyta) domoic/kainic acid pathways (Fig. 4). Notably, gene synteny for *dabA* and *dabC* is highly conserved among *N. navis-varingica* strains, with amino acid identities ranging from 84% to 99%. These findings underscore a unique evolutionary route for isodomoic acid B biosynthesis in *N. navis-varingica* and highlight the potential for divergent enzymatic adaptations in diatom versus red algal kainoid production.

**Figure 4.**
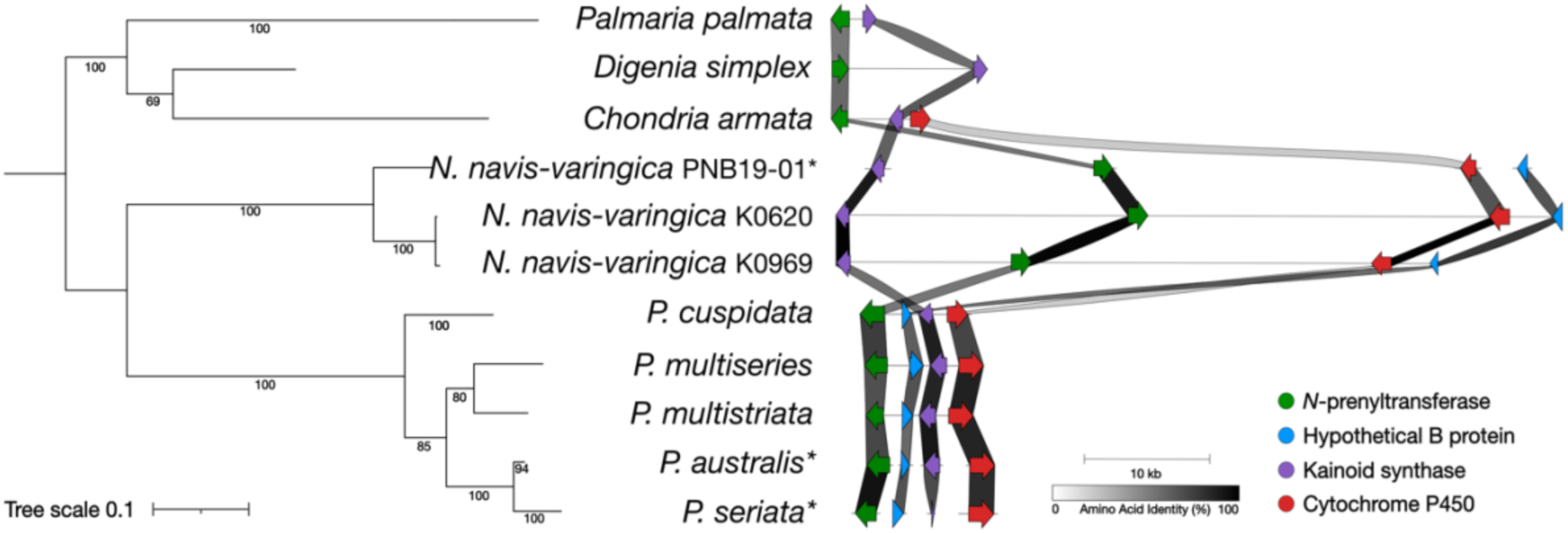
Syntenic comparisons and phylogenetic analysis of kainoid biosynthesis gene clusters. Phylogram represents unrooted phylogeny of concatenated kainoid biosynthesis genes coupled with visualization of the corresponding biosynthesis gene clusters. Lines represent contiguous stretches of DNA; arrows represent genes and their orientation; shading between genes represents amino acid sequence identity. *Transcripts.

## DISCUSSION

This study describes the DA biosynthesis pathway in pennate, benthic diatom *N. navis-varingica*. By combining genomic and transcriptomic analysis of regionally distinct strains of *N. navis-varingica*, we uncovered a novel organization of the *dab* gene cluster, assembled two high-quality diatom genomes, and clarified the phylogenetic position of *N. navis-varingica* within Bacillariaceae. Additionally, *in vitro* biochemical assays with key biosynthesis enzymes validated the DA isomer profile across strains, leading to the discovery of a diatom isodomoic acid B synthase, NnvDabC, which represents a novel enzymatic function within DA-producing diatoms.

### Gene cluster organization and evolution

The *dab* gene cluster in *N. navis-varingica* strains uniquely spans over 60 kb, which is roughly ten times larger than *dab* clusters found in *Pseudo-nitzschia* spp. and *C. armata* (15, 23). Expansion of BGCs through duplication and recruitment of additional enzymes has often been cited in eukaryotes for the diversification of secondary metabolite production (57–61). To our knowledge, this is the first instance of secondary metabolite BGC expansion in diatoms. The *dab* cluster expansion is further supported by an additional copy of *dabC* outside of the cluster, the co-localization of *dab* genes with repetitive elements, and interspersion of additional genes. Based on the genomic arrangement and gene synteny conservation among both *N. navis-varingica* strains and across diverse taxa, we hypothesize that core kainoid biosynthesis genes *dabA* and *dabC* form a single module within the cluster. The presence of flanking DNA transposons also supports the current proposal of DA evolution in which horizontal gene transfer contributed to the acquisition of this module. This observation is consistent with reports that diatoms may have acquired up to 5% of their genes through horizontal gene transfer (62). The downstream presence of *dabB* and *dabD* represent an additional module that was likely recruited through neofunctionalization and intra-genomic reorganization to confer the production of DA. The repetitive segment between *dabA* and *dabD*, displaying over 90% nucleotide identity in approximately 100 other places in the genome, further supports this hypothesis. Notably, protein of unknown function *dabB* is consistently found in diatom *dab* clusters but is absent in the red alga *C. armata*, suggesting a lineage-specific recruitment event that may contribute to differences in DA biosynthesis between these groups. Phylogenetic analysis of hypothetically assigned NnvDabD suggests its recruitment via neofunctionalization of native biochemistry, as was found in *C. armata* (15).

The conservation of a protein kinase downstream of the *dabBD* module—identified as a potential *dab* cluster integration site in *P. cuspidata*—suggests that this region may favor DA biosynthesis insertion and regulation (24). Phylogenetic analysis shows that the protein kinase is orthologous to sequences found in *P. multiseries* and *P. multistriata*, indicating that this locus may favor *dab* cluster insertion or facilitate genomic rearrangements involving the putative neofunctionalized CYP450. However, the CoA-binding gene present at the *dab* locus is not orthologous to sequences found in *Pseudo-nitzschia*; instead, the orthologous CoA-binding and protein kinase gene pair exists outside of the cluster in *N. navis-varingica*. This implies that their co-localization with the *dab* cluster may be coincidental, rather than representing a conserved regulatory or integration hotspot.

This regulatory context is particularly relevant when considering the evolution of the cluster toward DA production, which implies that DA confers ecological advantages, such as in response to pH or grazer stress (23, 26, 63–67). For example, the regulation of putative CYP450 DabD modulates the addition of the carboxylic acid moiety crucial for DA’s toxicity, supporting its potential role in deterring grazers (31). Indeed, the *N. navis-varingica* K0620 strain sequenced in this study was shown to deter grazing by the mixotrophic dinoflagellate *Karlodinium armiger* (68). Additionally, DA production may be linked to pH tolerance given that *N. navis-varingica* is found in shallow, brackish environments with elevated pH; whereas, DA production in *Pseudo-nitzschia* is linked with elevated pCO2 (23, 67, 69, 70). Further investigation of the regulatory network modulating DA production in response to environmental stress should be done by comparing gene expression patterns in *N. navis-varingica* and DA-producing *Pseudo-nitzschia* species.

### Mechanistic insights and enzymatic functions

Our work also sheds light on the evolution of the gene cluster towards DA production through functional diversification of kainoid synthase enzymes. Our biochemically supported phylogenetic analysis shows that isodomoic acid B synthase NnvDabC1 groups functionally with RadC, with both enzymes converting L-NGG (**3**) and cNGG (**4**) to DA-like molecules dainic acid A and isodomoic acid B (**5**), respectively. DA-producer *C. armata* RadC behaves similarly, whereas kainic acid-producing *D. simplex* KabC does not (15, 28). *P. multiseries* DabC was demonstrated to act as an isodomoic acid A synthase, whereas we have shown that *N. navis-varingica* DabC functions an isodomoic acid B synthase, demonstrating a distinct evolutionary trajectory between these genera.

The function of NnvDabC1, presence of additional genes at the *dab* locus, *dabC* paralogs outside of the *dab* cluster, and low homology of putative CYP450 NnvDabD, raises important biosynthesis questions about the order of operations in the pathway as well as the elusive final step—namely, the conversion of isodomoic acid A or B to DA. Conversion of isodomoic acid B to DA not only requires 1,3-olefin migration, but also an additional and separate *trans* to *cis* isomerization of the second olefin. Sequence analysis reveals that CYP450 NnvDabD exhibits significantly lower homology to other DabD enzymes in other *dab* and *rad* clusters, suggesting it may catalyze a distinct reaction. Consequently, rather than performing the precedented carboxylation of L-NGG (**3**), the pathway may proceed through the formation of dainic acid intermediates. This possibility is strengthened by the presence of dainic acids in culture extracts.

The discovery of additional oxidative enzymes—beyond the core alpha-ketoglutarate dependent Fe(II)-containing dioxygenase C protein and CYP450 D proteins—such as genes encoding acyl-CoA oxidase and aldehyde dehydrogenase within the *dab* locus may imply involvement in double bond isomerization. For example, these additional enzymes could induce oxidative rearrangement of the double bond to be shifted out of conjugation with the carboxylate, or work in concert with core oxidative enzymes in the *dab* cluster (71–73). Alternatively, though less precedented, activation of isodomoic acids by AMP-binding protein could allow for double bond isomerization by dehydrogenase enzyme, in a manner similar to bacillaene biosynthesis and other double bond isomerization biosynthesis strategies present polyketide systems (74–76). Although further work is required to fully elucidate the order and function of these steps, these potential mechanisms align with rare biosynthesis strategies in which oxidative enzyme-catalyzed saturation of unactivated double bonds results in olefin migration (72, 77, 78). However, the presence of *dabC* paralogs outside of the *dab* cluster implies that other genes outside of the *dab* locus may be involved in this pathway, as well.

### Ecological and evolutionary implications of genome expansion

Our study also found that *N. navis-varingic*a strains have exceptionally large genomes relative to other pennate and benthic diatoms. The over 750 Mbp genomes are more than fifteen times the size of closely related *N. inconspicua str. hildibrandii,* but only six times the size of fellow benthic diatom *Seminavis robusta* (35, 37). Similar to the observations of centric diatom *Thalassiosira* (with genomes up to 1.5 Gbp) and across eukaryotes, genome expansion in *N. navis-varingica* appears to be due to the expansion of noncoding DNA (43, 79, 80). This result is intriguing because Bergmann’s rule predicts smaller cells, and therefore genome sizes, at warmer temperatures, which are typical of sub/tropical benthic environments (81–83). Our findings add to emerging evidence that diatoms may defy this ecological principle, potentially contributing more to global primary production under the threat of rising ocean temperatures (84–86). In this instance, the size of *N. navis-varingica* genome may correlate with increased genetic diversity and adaptations to fluctuating salinity, pH, light, and nutrient availability, which in turn could lead to greater cell abundance—a trend in support of the latter hypothesis and observed in polar diatom communities (43). Moreover, DA-producing species of *Pseudo-nitzschia* have augmented genomes with increased repetitive elements, relative to non-DA producing species (24, 25). Genomes of non-DA producing strains of *N. navis-varingica* may reveal a similar trend of differential DNA content.

### Taxonomic and phylogenomic considerations

Following the first outbreak of DA poisoning by the diatom formerly known as *Nitzschia f. pungens* (now, *Pseudo-nitzschia multiseries*), the genus *Pseudo-nitzschia* was delineated from *Nitzschia* partly through taxonomic research driven by DA production (6, 7). Our work extends this re-evaluation to *N. navis-varingica*, another DA-producer within the Bacilliaracae family, which has been documented as widespread throughout the Western Pacific and harbor significant intraspecific diversity (22, 34). Our analysis of multiple conserved genetic markers of the speciose *Nitzschia* genus and Bacillariaceae family reveals that *N. navis-varingica* strains clade with a subset of *Nitzschia* spp. and *Psammodictyon* spp. commonly found in the benthos of the western sub/tropical Pacific. Given its phylogenetic position outside the primary *Nitzschia* clades, the reported absence of unifying synapomorphy, and presence of DA-producing species, this “cryptic” clade is a strong candidate for taxonomic revision (34, 87). Further phylogenomic investigations are recommended to resolve these taxonomic ambiguities and elucidate the origins of DA production within the Bacillariaceae.

## Conclusion

In summary, our study provides new insights into the evolution and organization of the DA biosynthesis pathway by identification of a novel *dab* cluster in *N. navis-varingica*. The unique genome-wide expansion reflects the expansion and modular reorganization of the *dab* gene cluster. The discovery of the genetic basis of DA production in *N. navis-varingica* and biochemical verification of the Dab pathway suggest that DA production may offer ecological advantages in the subtropical benthic habitat. Future research should explore *N. navis-varingica dab* gene expression patterns and taxonomic relationships among DA-producing diatoms to better understand the ecological and evolutionary implications of DA production.

## MATERIALS AND METHODS

### Diatom culturing and harvesting

*Nitzschia navis-varingica* strains K0620 and K0969 were obtained from the Norwegian Culture Collection of Algae (NORCCA, https://norcca.scrol.net/, 39). Strains were maintained in natural seawater F/2 media (Guillard, 1983), 16°C, and under a 12:12 photoperiod. Strain K0969 was incidentally co-cultured with a *Pseudobodo* sp. (Bicoecea), present from the culture collection. Attempts to isolate strain K0969 without the contaminant were unsuccessful.

PNB19-01 (19) was cultured in 30 mL of F/2 medium in 50 mL tissue culture flask (Greiner bio-one, Tokyo, Japan), and by incubating them at 25°C under an irradiance level of 80 μmol photons m^−2^ s^−1^, with a 12:12 h light:dark cycle. The medium was prepared using seawater diluted with distilled water to a salinity of ca. 28.

For scale-up culturing, K0620 and K0969 were first treated with antibiotics. They were then grown at 4 L scale at 20°C until exponential growth phase, approximately 1 week. To maintain exponential phase, 2 L of cell culture was harvested by centrifugation at 7,000 xg for 20 minutes every 4-5 days. The remainder of the culture was replenished with 2 L F media. This was repeated until sufficient biomass was collected.

### DNA and RNA extraction

For strains K0620 and K0960, high molecular weight (HMW) DNA was extracted from 0.1 g biomass using Illustra Nucleon Phytopure Genomic DNA Extraction Kit (Cytiva). Manufacturer’s instructions were followed with the following exception: the chloroform extraction and DNA precipitation steps were repeated 5 times in order to increase the quality of DNA. Extracted DNA was size selected using the BluePippin system with a High Pass Plus 15 kb cassette (Sage Science Cat# BPLUS03) and HMW fragment lengths verified using the 4150 TapeStation (Agilent Cat# G2992AA) with a Genomic DNA Screentape (Agilent Cat# 5067-5365). RNA was extracted from 0.1 g of biomass using the Direct-zol RNA Purification Kit (Zymo) following the manufacturer instructions. RNA was quantified and quality assessed using a Qubit RNA BR Assay Kit (Invitrogen Cat# Q33231) and TapeStation RNA ScreenTape (Agilent Cat# 5067-5576).

For strain PNB19-01, the 29 days culture (15 mL) was harvested at 3.3 h after switching the lighting to light from dark by centrifugation in 50 mL conical tube at 670g for 1 min at 25°C. After removal of the supernatant, TRI reagent (Sigma, cat#T9424, 1 mL) and 0.1 mL of Zirconia Ball YTZ-0.05 mm (Nikkato corporation, Japan) were added to the cells, then the cells were disrupted using MS-100 (TOMY, Japan), 3,000 rpm for 1 min. The suspension was moved into a micro tube (1.5 mL) and chloroform (0.2 mL) was added, then kept for 5 min at room temperature (RT). The mixture was centrifuged for 15 min at 20,600g at 4°C. The supernatant was moved to a new microtube, then isopropanol (0.5 mL) was added and kept for 5 min at room temperature (RT). After centrifugation for 15 min at 20,600g at 4°C, the supernatant was removed, then 75 % EtOH (1 mL) was added to the precipitation, then centrifuged again for 6 min at 5,000g at 4°C. After removal of the supernatant, the precipitation was dissolved with RNase free water (15 µL), and mixed with DNase I 10 x buffer (1.5 µL) and DNase I (RNase free, Nippon gene, 0.5 µL) using vortex, then kept at 37°C for 10 min. After reaction, TRI reagent (0.5 mL) and chloroform (0.1 mL) were added to the reactant and kept 5 min at RT, then centrifuged for 15 min at 20,600g at 4°C. The supernatant was moved to a new microtube, then isopropanol (0.25 mL) was added. After keeping at RT for 5 min, the mixture was centrifuged for 15 min at 20,600g at 4°C. The supernatant was removed, then 75%EtOH (0.5 mL) was added, then centrifuged 6 min at 5,000g at 4°C. The obtained total RNA was dissolved with 50 µL of RNase free water, then quantified as total 1.38 µg using Quantus Fluorometer (Promega).

### Library preparation and sequencing

Strains K0620 and K0969 were sequenced on a Pacific Biosciences (PacBio) Revio to produce PacBio HiFi reads, intended for nuclear genome assembly, and IsoSeq reads were used to map transcriptomic data to the assembled genome. SMRTbell libraries were prepared from the HMW DNA preps of each strain using the HiFi SMRTbell prep kit 3.0 (PacBio Cat# 102-182-700) according to the manufacturer’s instructions, including the recommended DNA shearing step for eukaryotes. Iso-Seq Kinnex libraries were prepared from total RNA using the Kinnex Full-Length RNA kit (PacBio Cat# 103-238-700) with the Iso-Seq Express 2.0 Kit for cDNA synthesis (PacBio Cat# 103-071-500), according to the manufacturer’s instructions which include a polyA-selection. The K0620 and K0969 SMRTbell libraries were barcoded and multiplexed on a single 25M SMRT cell (Cat# 102-202-200), and the K0620 and K0969 Kinnex libraries barcoded and multiplexed on a single SMRT cell along with several other libraries. Both runs used the v13.0.0.205983 Revio chemistry bundle and a 30-hour movie time.

For strain PNB19-01, cDNA libraries were sequenced by Novogene using NovaSeq X Plus (Illumina) (3Gb, PE150, 20 M PE read).

### Genome and transcriptome assembly, size estimation, and annotation

The PacBio genomic HiFi reads from the *N. navis-varingica* K0620 and K0969 samples were assembled into partially phased contigs using HiFiasm v0.19.8. The assembled contigs were then screened and filtered for contamination utilizing v0.5.0 of NCBI’s Foreign Contamination Screening – GX (FCS-GX) workflow. The filtered assemblies were then assessed for contiguity and completeness with assembly-stats v1.0.1 and the stramenopiles_odb10 BUSCO v5.4.3 database.

Concurrently, a k-mer approach was taken to predict genome size, heterozygosity, and repeat content of the *N. navis-varingica* samples using the HiFi reads, GenomeScope 2.0, and meryl v1.3. For sample K0969, lower HiFi read coverage required setting the initial kmercov estimate to 10 for GenomeScope 2.0.

Transcripts were identified from the PacBio IsoSeq reads of *N. navis-varingica* K0620 and K0969 using isoseq v4.2.0. The reads were segmented with skera v1.3.0 before primer removal and read demultiplexing via lima v2.12.0. After segmentation and demultiplexing, poly(A) tails and concatemers were removed with the “isoseq refine --require-polya” command. The three IsoSeq runs were then clustered for each sample by running “isoseq cluster2 --singletons” and subsequently mapped to the associated genome assembly with pbmm2 v1.16.0.

Genome annotation was conducted using a combination of BRAKER v2.1.6, GALBA v1.0.1, and TSEBRA v1.1.2.5. BRAKER was used to predict genes based on the IsoSeq transcript mapping and protein annotation sequences from 8 related Bacillariophyceae species: *Phaeodactylum tricornutum CCAP 1055/1, Fragilariopsis cylindrus CCMP1102, Fistulifera solaris, Nitzschia inconspicua, Mayamaea pseudoterrestris, Pseudo-nitzschia multistriata, Seminavis robusta,* and *Cylindrotheca closterium* with the GenBank accessions: GCA_000150955.2, GCA_001750085.1, GCA_002217885.1, GCA_019154785.2, GCA_027923505.1, GCA_900660405.1, GCA_903772945.1, and GCA_933822405.4, respectively. These protein sequences were additionally used in the GALBA annotation workflow to predict protein coding gene structures. After BRAKER and GALBA, the results were input into TSEBRA to select the highest-confidence transcripts. Functional annotations of gene models were generated by annotation against common protein domain databases Pfam v35.0, PANTHER v15.0, TIGRFAM v15.0, KEGG v30-01-2023, and EggNOG v5 using a combination of tools (diamond v2.0.15, eggNOG-mapper v2.1.10, Interproscan v5.57-90.0, kofamscan v1.3.0 (88–93). Lineage Probability Index (LPI) was calculated from top 100 diamond blastp hits by dividing the sum of probabilities of each taxonomic term by the normalization factor corresponding to its taxonomic level in the lineage and choosing the term with the highest index (94). For each assembly, RepeatModeler v2.0.6 with default parameters was used to prepare a custom repeat library which was used as input for RepeatMasker v4.1.8 with the “-xsmall -nolow -norna -no_is -q” parameters.

For PNB19-01, *de novo* assembled data was used for BLAST search and other bioinformatics analysis using GENEYX-MAC (Nihon Server, Tokyo).

### Annotation of domoic acid biosynthesis gene clusters and phylogenetic analysis

*Dab* gene amino acid sequences from *P. multiseries*, *P. multistriata*, *P. australis*, and *C. armata* were used to build a Hidden Markov Model (HMMER v3.3.2) query for each individual gene in the cluster using hmmsearch (part of HMMER) or BlastP (95–97). The peptide sequences from assemblies were queried to identify candidates and verified by sequence alignment.

Single gene phylogenies were built using Kalign (EMBL-EBI), top BLAST hits for the *N. navis-varingica* sequences, representative UniRef50 sequences, in addition to those listed in previously published phylogenies (15, 98). Concatenated phylogenies of Bacillariaceae marker genes (Fig. 1) and kainoid biosynthesis genes (Fig. 4) were built using SPLACE to align and concatenate genes of interest (99). Maximum-likelihood trees were built using IQ-TREE and visualized in iTOL (90, 91).

Visualization of genomic data was performed using RStudio (v2024.12.1.563), including packages taxize, gggenes and ggplot2 (100–103). Clinker was used for kainoid gene cluster comparison (104). Subsequent figure refinement (color adjustments and figure compilation) was performed in Affinity Designer (https://affinity.serif.com/en-us/designer/).

### Heterologous protein expression and purification

For strain PNB19-01, *dabA* and *dabC* clones were obtained using reverse transcriptase polymerase chain reaction (RT-PCR). They were then cloned into a pet28 vector and expressed as described below. See all expressed genes in Table S3. In RT-PCR obtained clone of *dabA*, single nucleotide change from 296-A (RNA-seq) to G was detected. This 296-G clone was expressed.

Putative chloroplast signal peptide was identified on K0620-dabA using HECTAR (v1.3) and SignalP (Eukarya, v6.0) (55, 56). Therefore, K0620 dabA was ordered with a 26 amino acid truncation and expressed as such. Both *dabA* and *dabC* genes were codon optimized for *E. coli* expression, domesticated for SapI and BsaI cut sites, and designed with N-terminal His-6 affinity tag in pET-28a(+) vectors and ordered from Twist Bioscience. See all expressed nucleotide sequences in Table S3.

Both DabA and DabC were expressed and purified as previously (15, 23) using conventional methods. Constructs were transformed into chemically competent *E. coli* BL21(DE3) cells and plated on kanamycin (50 mg/mL) plates. Overnight cultures of transformed BL21(DE3) *E. coli* were used to inoculate expression cultures, which were grown at 37°C in 500 mL TB broth supplemented with 4% glycerol to an OD-600 of ∼0.6. Cultures were chilled on ice and induced with 1 mM of isopropylthio-beta-galactoside (IPTG). Flasks were shaken at 18°C overnight (∼18 hours). Cells were harvested by centrifugation (8,000 x g, 15 mins) and frozen at -80°C until future purification.

Frozen pellets were defrosted on ice and at 4°C overnight, resuspended in 5 mL lysis buffer (10 mM HEPES, 100 mM NaCl, 25 mM Imidazole, 0.2 mM DTT, 2.5 mM EDTA, 20% glycerol, pH 7.5) per 5 mg pellet amended with 1 mg/mL lysozyme. Cells were lysed by sonication using a Qsonica tip at 50% amplitude for 15s on, 45s off, 7 min total working time. DNase I in 5 mM MgCl2 was added halfway through sonication to a final concentration of 5 ug/mL. Lysate was centrifuged at 40,000xg for 30 minutes at 4°C to remove cellular debris. Supernatant filtered through Whatman filter before purification.

Purification was performed using immobilized metal-affinity chromatography purification (IMAC) of His6-tagged proteins using a HisTrap FF column (Cytiva) on a AKTA pure™ 25 L1 (Cytiva) fast protein liquid chromatography (FPLC) system and a BioLogic DuoFlow system (Bio-Rad). FPLC data was analyzed with UNICORN version 7 software. Clarified lysate was loaded at 2 mL/min onto a 5 mL HisTrap FF column (Cytiva) pre-equilibrated with lysis buffer. The column was washed with 10 column volumes of 8% elution buffer (10 mM HEPES, 100 mM NaCl, 500 mM imidazole, 20% glycerol, pH 7.5) and then eluted with a linear 8–100% gradient over 15 column volumes in 4 mL fractions. Fractions were analyzed by SDS-PAGE. Fractions containing the target protein were pooled, desalted, and buffer-exchanged using PD-10 columns (Sephadex G-25 M, Cytiva) that were pre-equilibrated with storage buffer (50 mM HEPES, 250 mM NaCl, pH 8, 10% glycerol). Storage buffer for DabA was amended to final concentration of 5 mM MgCl2.

Buffer exchanged protein was concentrated using Amicon Ultra-15 centrifugal filters, aliquoted, and flash frozen in liquid nitrogen. Aliquots were stored at -80 until further analysis. All protein quantification was calculated using denatured protein UV absorbance at 280 nm and the protein’s extinction coefficient, at multiple dilutions with Milli-Q water.

### Preparation of substrates and standards and enzymatic activity assays

DA standard was purchased from the National Research Council of Canada (105). Kainic acid standard was purchased from Chem-Impex. Preparation of all non-commercial substrates were used as prepared for previous studies (15, 23, 28, 31). All substrates and enzymatic assay products were verified using retention time and mass via LCMS.

DabA enzyme assays to demonstrate N-prenyltransferase function were performed as previously described (15, 23, 30), with a few modifications. Reaction mixture was prepared in a final volume of 100 μL in 100 mM HEPES (pH 8.0), 100 mM KCl, 10% glycerol buffer with 5 mM MgCl2, 1 mM geranyl diphosphate (GPP), and 20 mM of L-glutamate. Reactions were allowed to incubate at room temperature (∼22 °C) for 6 hours and were then quenched with 100 L of ice-cold methanol. Quenched reactions were centrifuged, filtered, and injected (10 uL) onto LC-HRMS.

Enzyme assays to demonstrate kainoid synthase activity for K0620-DabC and PNB19-01-DabC were performed as previously described (15, 23) with few modifications. A reaction mixture was prepared in a final volume of 100 μL containing 100 mM HEPES (pH 8.0), 100 mM KCl, 10% glycerol, 1 mM L-ascorbate, 6.25 mM 2-oxoglutaric acid, 25 μM to 1 mM of either 7’-COOH-L-NGG or L-NGG, 50 μM DabC, and 50 μM FeSO₄·7H₂O, and incubated at 25°C overnight (15 h). The reaction was quenched by adding 100 μL of methanol, followed by centrifugation. The supernatant was purified via filtration with a nylon 0.22 µm pore CA membrane (Costar Spin-X) or purified using a reversed-phase resin (Cosmosil 140C-OPN) prior to liquid chromatography mass spectrometry (LCMS) analysis.

### Metabolite extraction from *N. navis-varingica*

Solid phase extraction was performed using Agilent Bond Elut PPL cartridges (200 mg, 3 mL), in a manner similar to analysis for dissolved organic matter analysis (106). During exponential phase of strains K0620 and K0969, 50 mL of cell culture was acidified to pH 2 with concentrated HCl. The PPL cartridges were washed and activated using LCMS grade methanol and LCMS grade water (pH 2), respectively. Acidified samples were loaded onto the cartridge under vacuum in a drip-wise manner. After sample loading, cartridges were washed with acidified LCMS grade water (pH 2) to remove salts. Cartridges were dried under nitrogen gas and eluted with 2 mL of LCMS grade methanol. Samples were dried down in a Speedvac and resuspended in 100 µL of 80% aqueous methanol (LCMS grade) with 0.1% formic acid (LCMS grade). Samples were stored at -80°C until LCMS analysis.

PNB19-01 cell extract (the one-month culture, 8 mL) cells (approximately 10 mg) were collected by 1,500 × g 3 min centrifugation, then extracted with 50% MeOH 100 µL by sonication 10 sec. After centrifugation (20,600 × g 30 sec), the supernatant was collected. The half of the supernatant (approximately 50 µL) was used after removal of the solvent using vacuum centrifugation at room temperature (finally from 5 mg cell extract).

### Liquid chromatography mass spectrometry

LCMS measurements were conducted in a similar manner as previously described (11). Samples were injected onto an Agilent single quadrupole UPLC-MS iQ using the Single Quadrupole Analytical LCMS. Compounds were separated by reversed-phase chromatography on a Phenomenex Kinetex 5 mm C18 100 Å 150 x 4.6 mm LC column with water + 0.1% formic acid (solvent A) and acetonitrile + 0.1% formic acid (solvent B) as eluents. The following gradient was applied at a flow rate of 0.75 mL/min: hold at 5% B for 1 minute, 5% to 35% B over 30 min, 35 to 100% B over 1 minute, hold at 100% B for 1.5 min, 100% to 5% B over 2.5 min, hold at 5% B for 2 min.

Higher resolution mass spectrometry was necessary for detection of biosynthetic intermediates in culture extracts. High resolution LCMS measurements were conducted using an Agilent Technologies 1200 Series system with diode array detector coupled to an Agilent Technologies 6530 accurate-mass Q-TOF LCMS. Identical chromatographic methods were applied and run in negative ionization mode.

## ACKNOWLEDGMENTS

This work was supported by the National Oceanic and Atmospheric Administration (NA19NOS4780181 to B.S.M. and A.E.A.), JSPS KAKENHI (JP23H02146 and JP23K26839 to M.Y.Y.), Tang Genomics Fund (T.P.M.), and graduate student fellowships from UC San Diego (S.M.W.-R., A.B.) and the National Science Foundation (NSF) (GRFP #2021321499 to M.L.M.).

---

**Figure S1.**
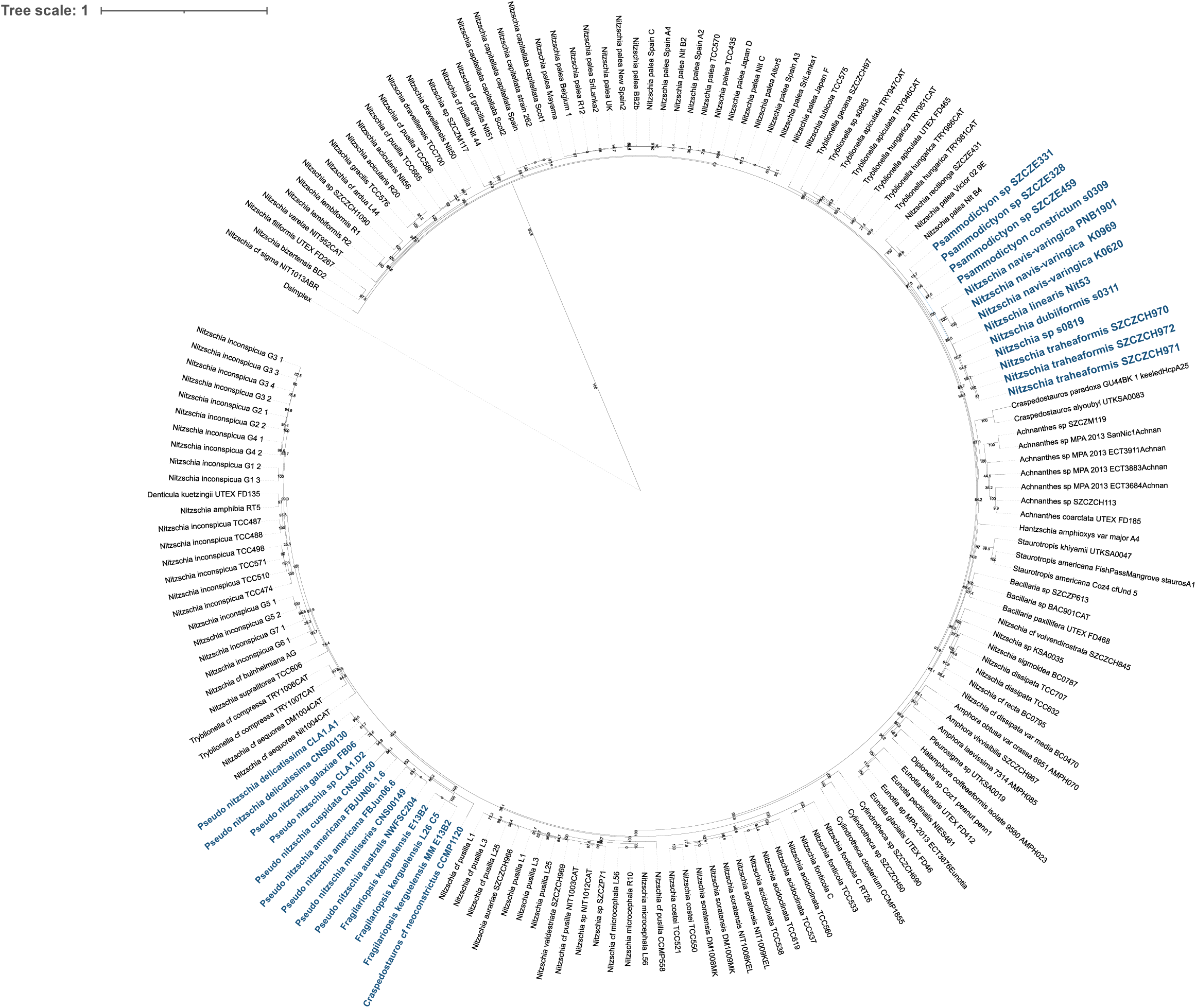
Rooted phylogenetic analysis of Bacillariaceae family using concatenated alignments of 18S (SSU) and 23S (LSU) nuclear RNA genes as well as chloroplast genes psbC and rbcL. *Digenea simplex* genes used as outgroup. Blue clades are included in Fig.1. Bolded are strains highlighted in this study.

**Figure S2.**
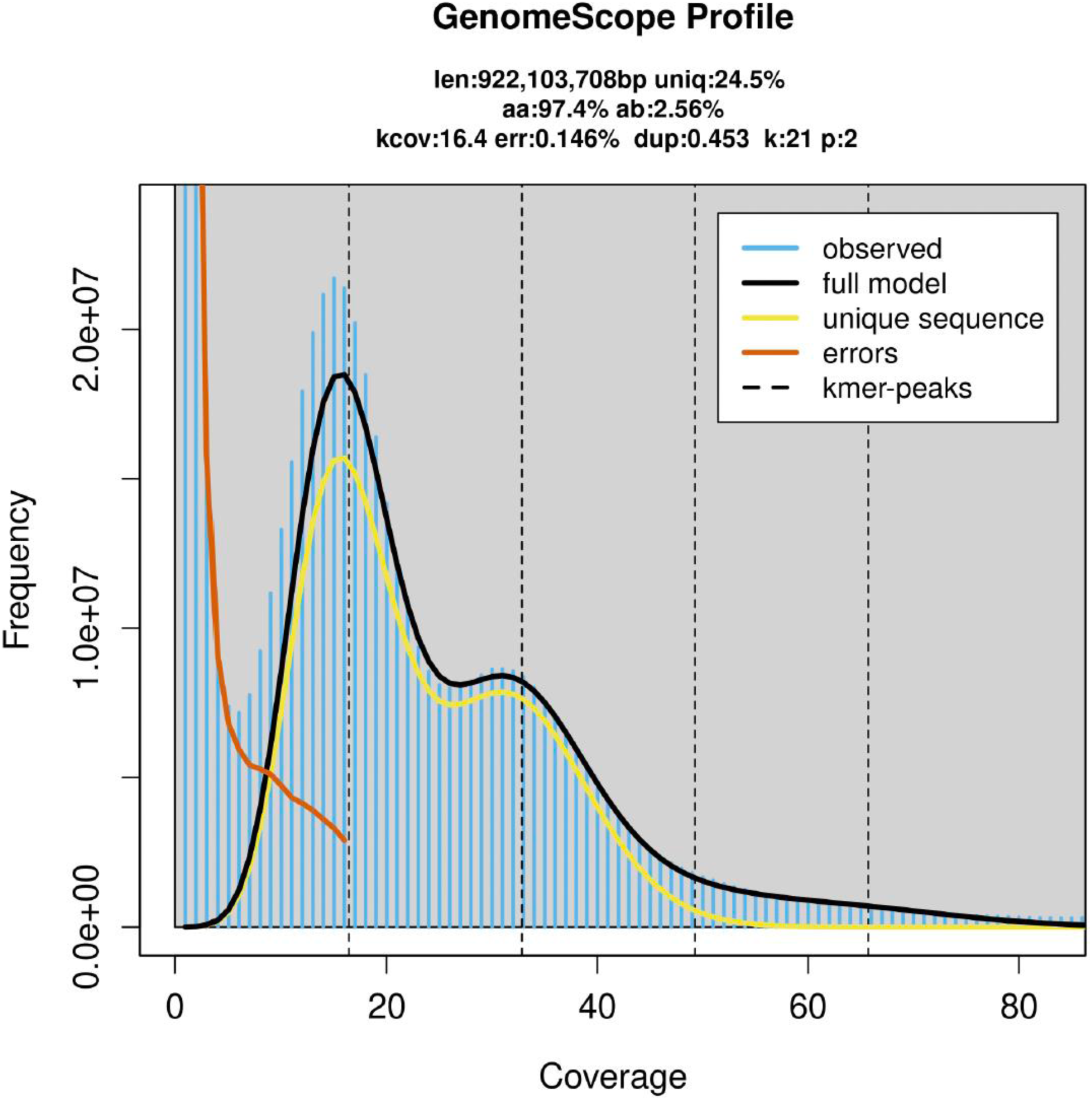
Genome survey analysis of *N. navis-varingica* K0620. _30_

**Table S1.** Genome statistics for assemblies produced for *N. navis-varingica* K0620 and K0969.

| Assembly | K0620<br>consensus<br>(v1) | K0620 hap1 | K0620 hap2 | K0969<br>consensus (v1) | K0969 hap1 | K0969 hap2 |
| --- | --- | --- | --- | --- | --- | --- |
| Size (bp) | 1,140,675,705 | 879,516,262 | 719,094,303 | 1,047,029,800 | 782,847,849 | 624,816,545 |
| Number of contigs | 4,911 | 5,895 | 3,355 | 8,655 | 9,152 | 6,055 |
| N50 (bp) | 442,841 | 303,884 | 342,993 | 198,174 | 134,724 | 152,664 |
| BUSCO Complete (%) | 99 | 94 | 92 | 96 | 86 | 87 |
| BUSCO Duplication (%) | 43 | 19 | 20 | 55 | 31 | 17 |
| GC content (%) | 31.75 | 31.95 | 31.24 | 33.37 | 33.89 | 33.77 |
| Repeat region (%) | - | 71 | 70 | - | 70 | 71 |
| Coding sequences (%) | - | 7.17 | 7.15 | - | 7.12 | 7.21 |

**Figure S3.**
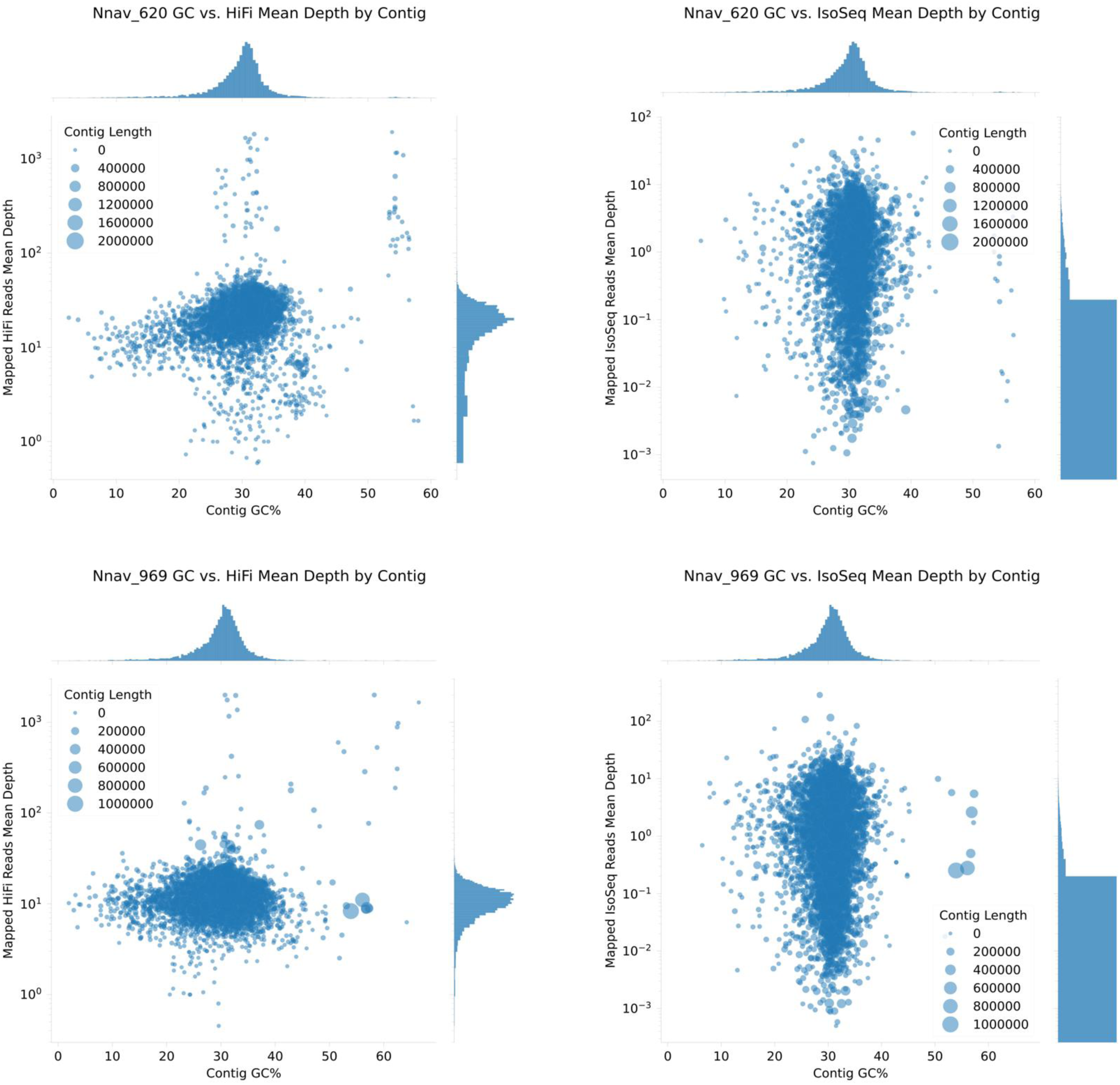
Mean Depth of IsoSeq and HiFi reads to Hap1 of Nnav620 and Nnav969.

**Figure S4.**
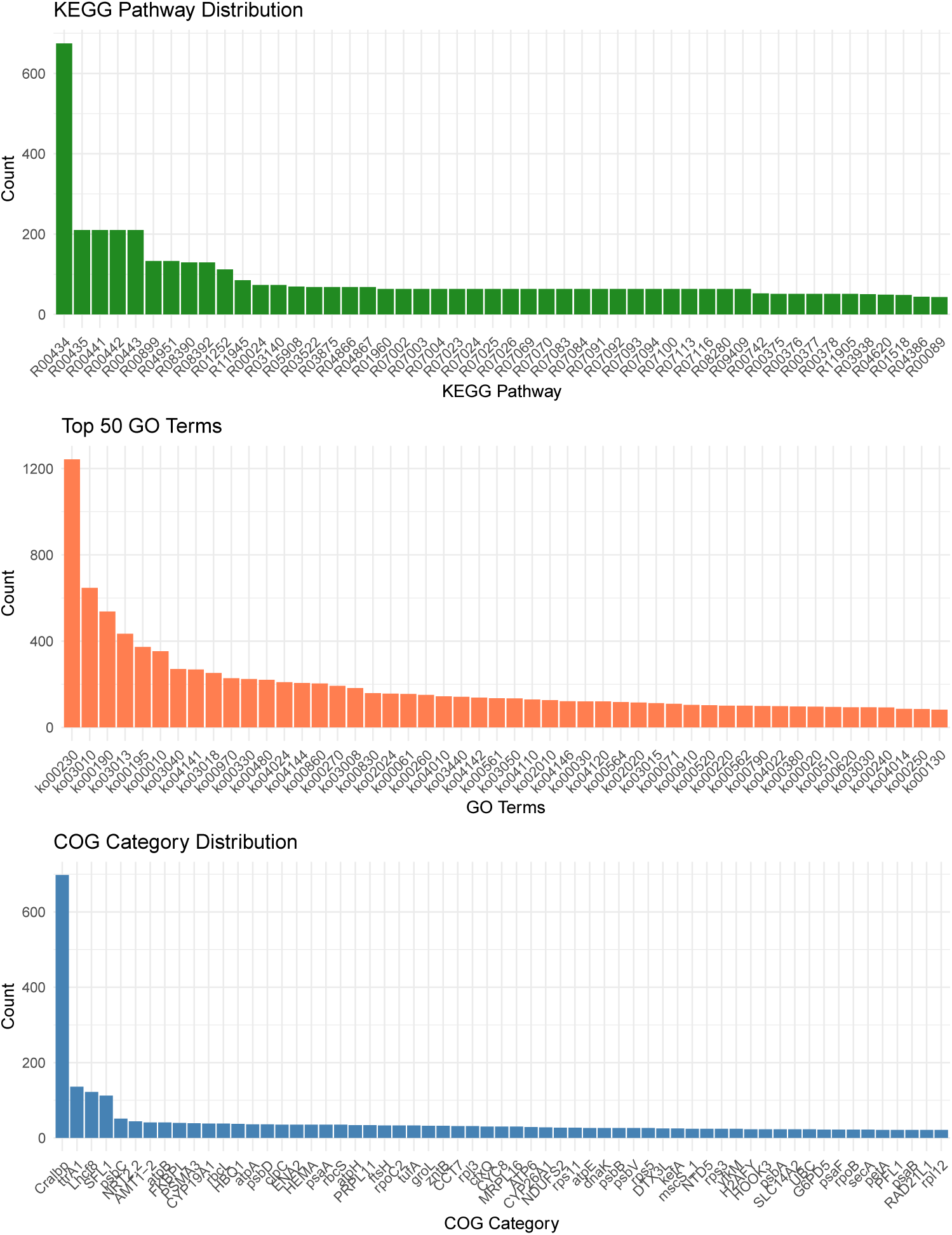
Distribution of KEGG, GO, and COG terms in EggNOG mapper functional annotations. 33

**Table S2.**
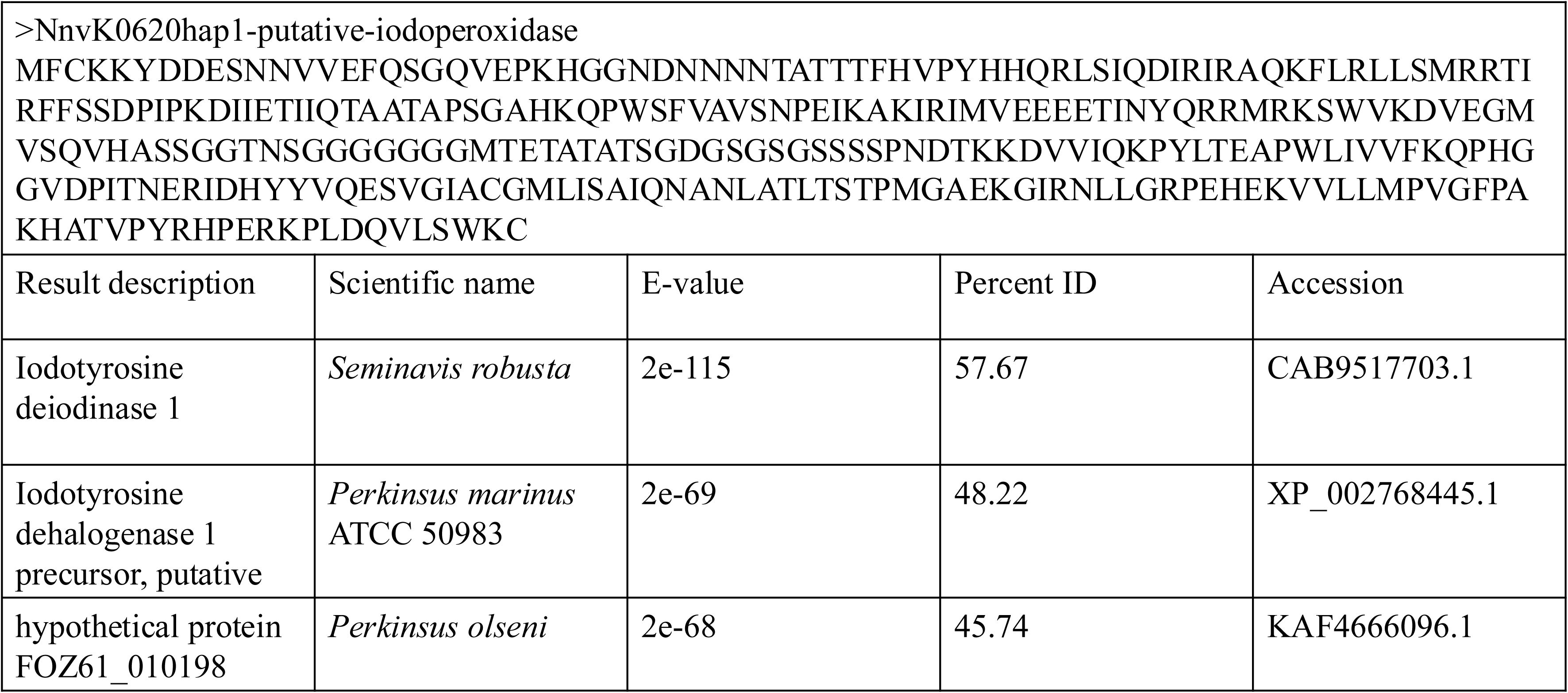
Amino acid sequence for putative iodoperoxidase gene from K0620 hap1 assembly. Top BlastP hits from NR database.

**Figure S5.**
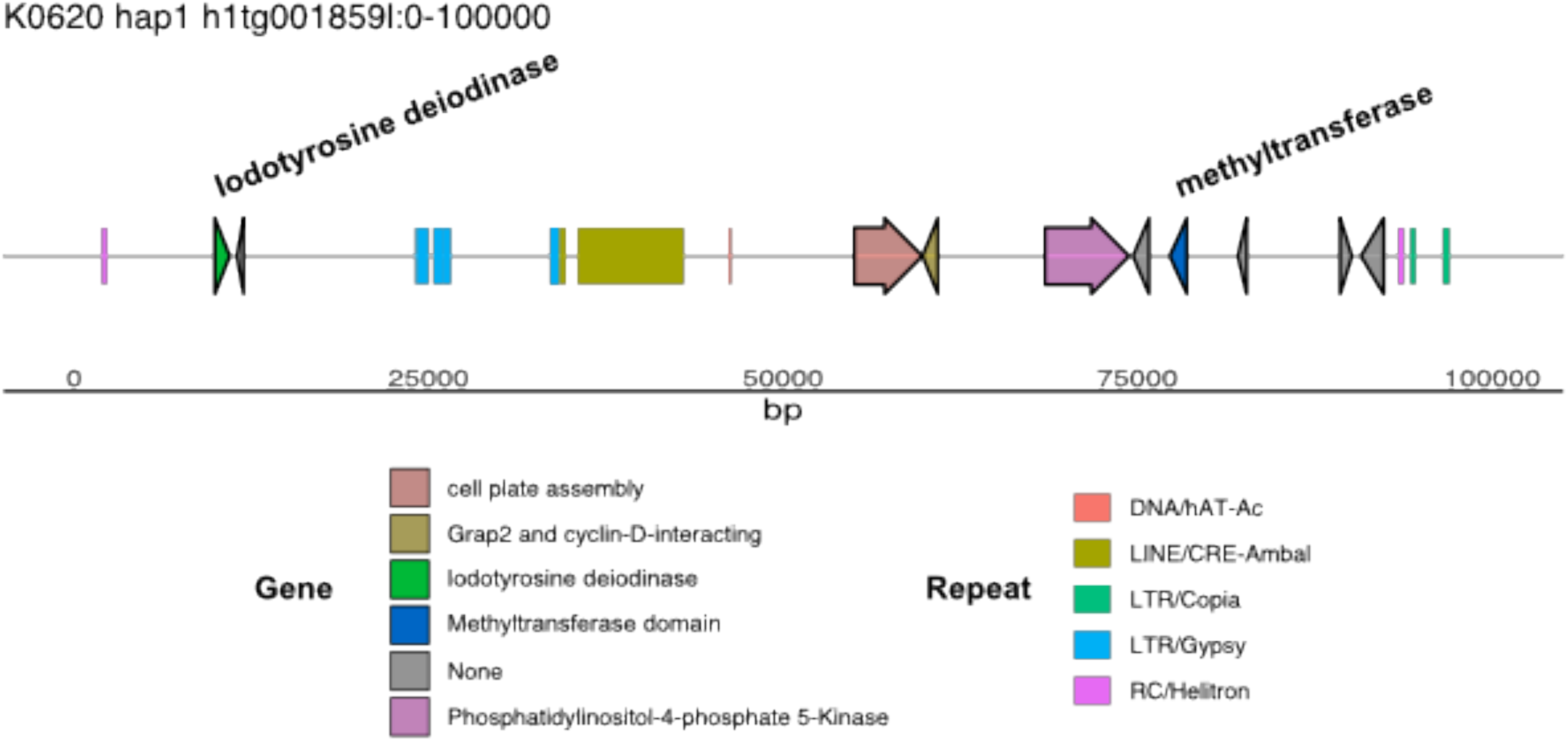
Putative iodotyrosine deiodinase gene containing contig in K0620 hap1. Arrows represent genes and bars represent repetitive elements.

**Figure S6.**
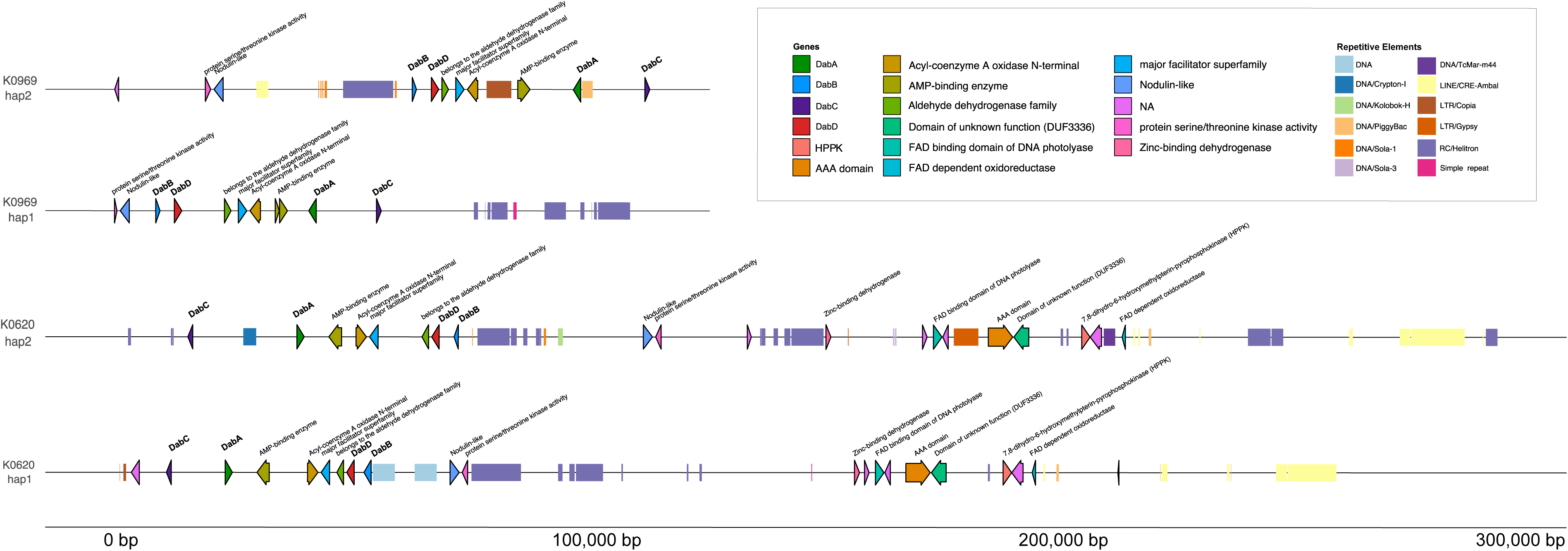
Domoic acid biosynthetic gene cluster containing contigs across both haplotype-resolved assemblies for strains K0620 and K0969. Arrows represent genes and bars represent repetitive elements.

**Figure S7.**
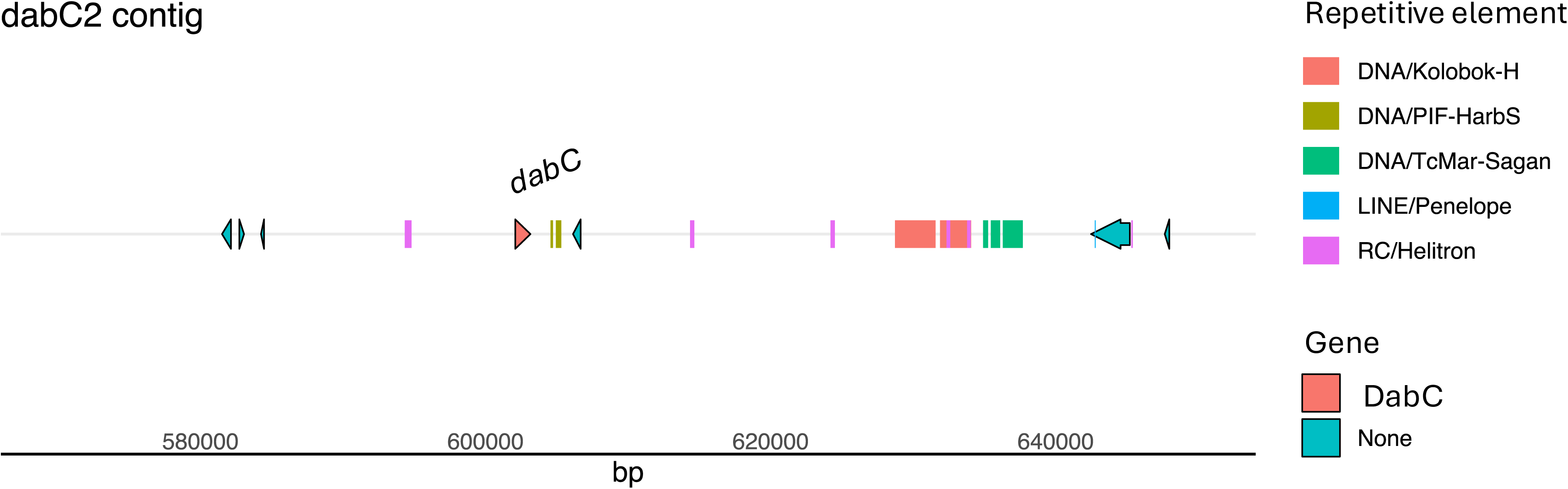
*N. navis-varingica* hap1 *dabC2* contig genomic neighborhood. Arrows represent genes and bars represent repetitive elements.

**Fig S8.**
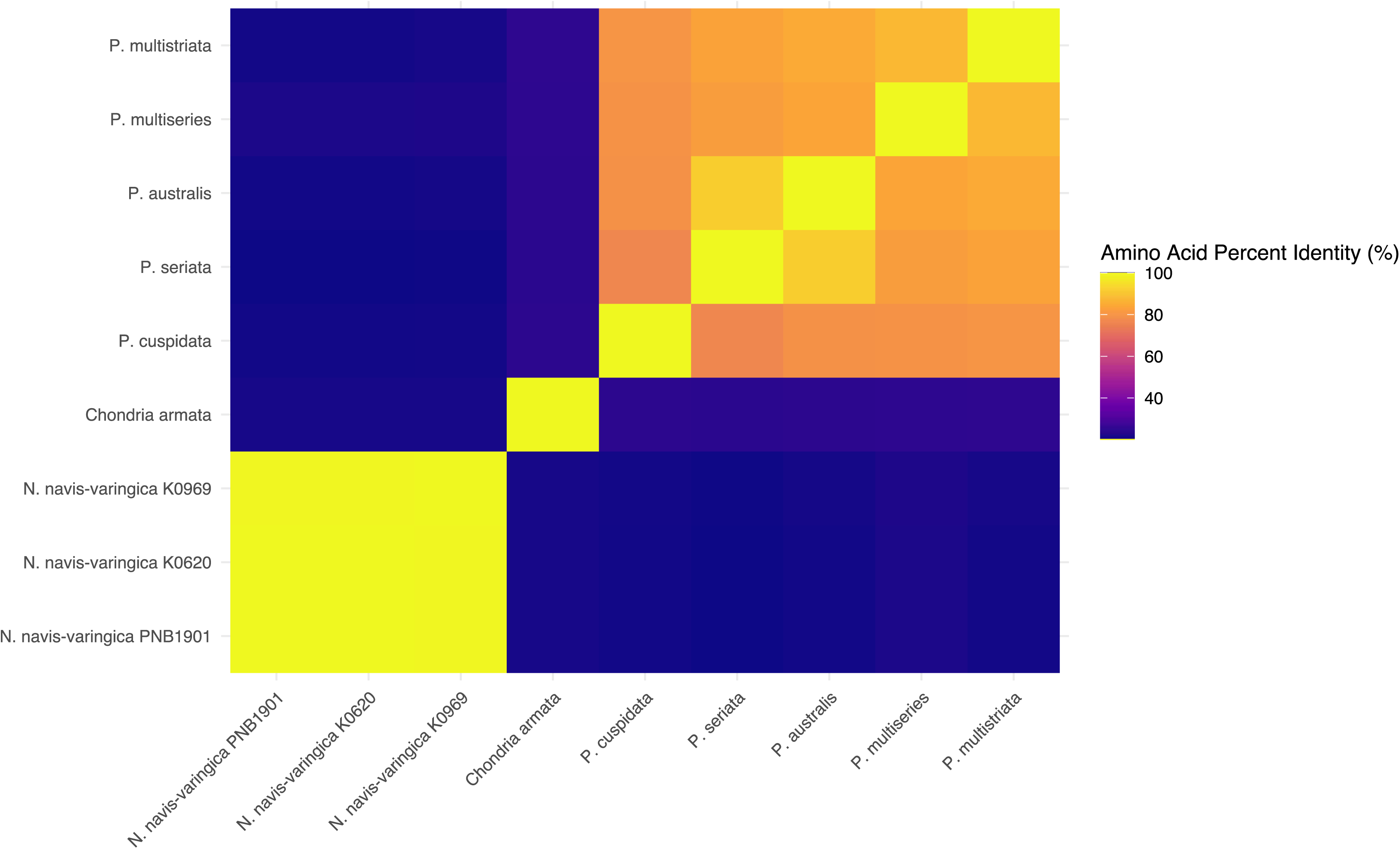
Heatmap of amino acid percent identity among dabD sequences.

**Figure S9.**
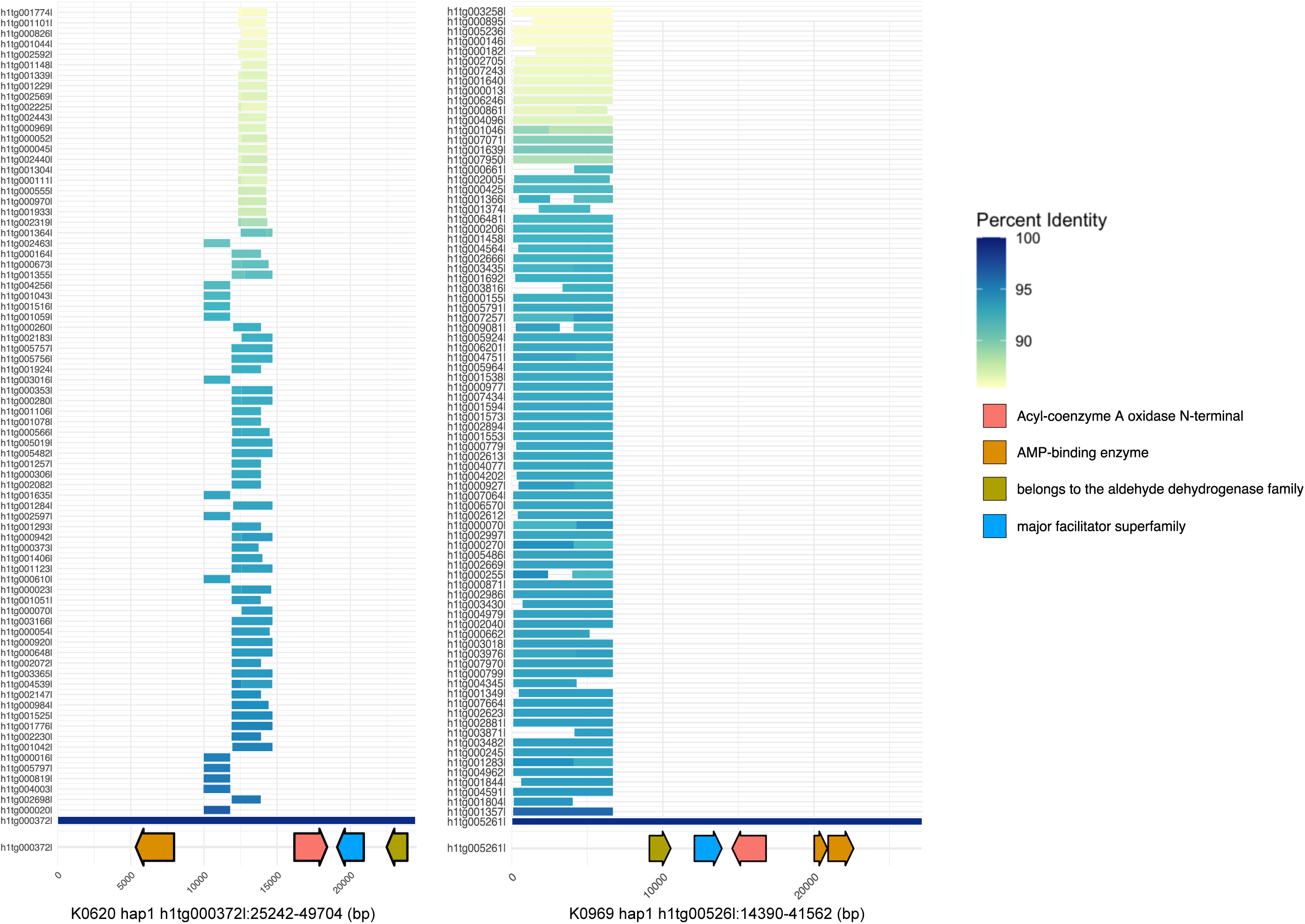
Comparative analysis of top 100 BLAST hits for *dab* cluster segment repeat element in K0620 hap1 and K0969 hap1. Each row represents a contig, horizontal bars show the aligned portion of the segment repeat and indicate nucleotide percent identity (85%+) relative to the reference *dab*-containing contigs shown on the bottom row. The schematic at the bottom illustrates the portion of the *dab* cluster between *dabA* and *dabD,* with coding sequences represented as arrows.

**Figure S10.**
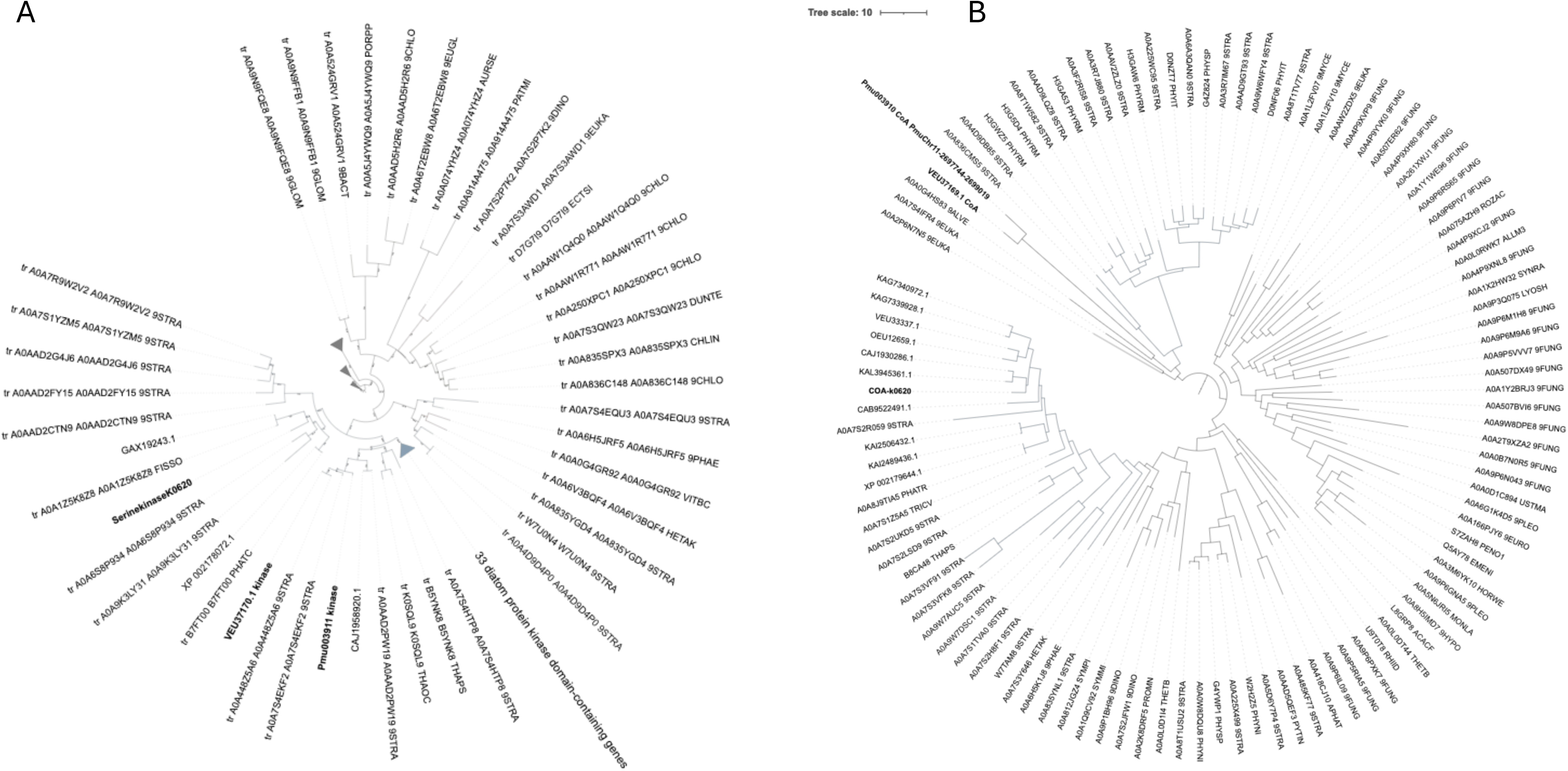
Phylogenetic analysis of (A) protein kinase and (B) CoA-binding protein genes. *Nitzschia navis-varingica* and *Pseudo-nitzschia* spp. genes are bolded. Blue branches correspond with diatom sequences.

**Figure S11.**
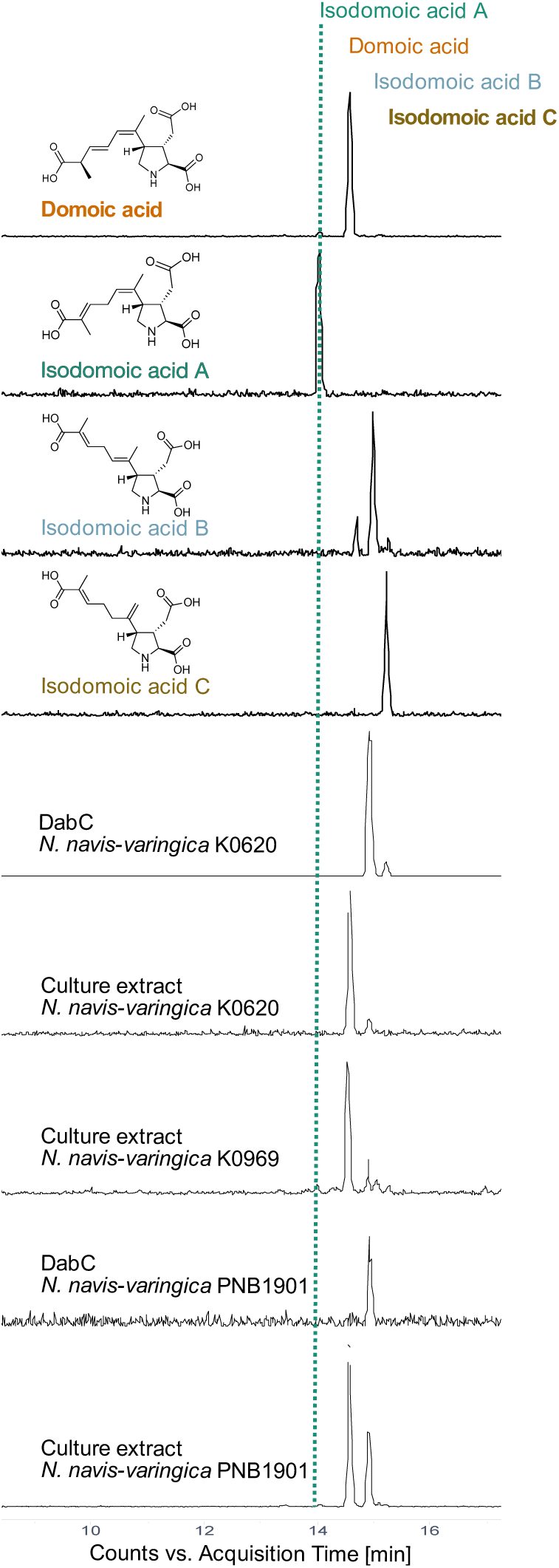
Culture extracts and DabC assays compared with synthetic standards. Positive ion mode LC-MS extracted ion chromatogram profiles for anticipated domoic acid and related isomer products (*m/z* 312.1±1.0).

**Figure S12.**
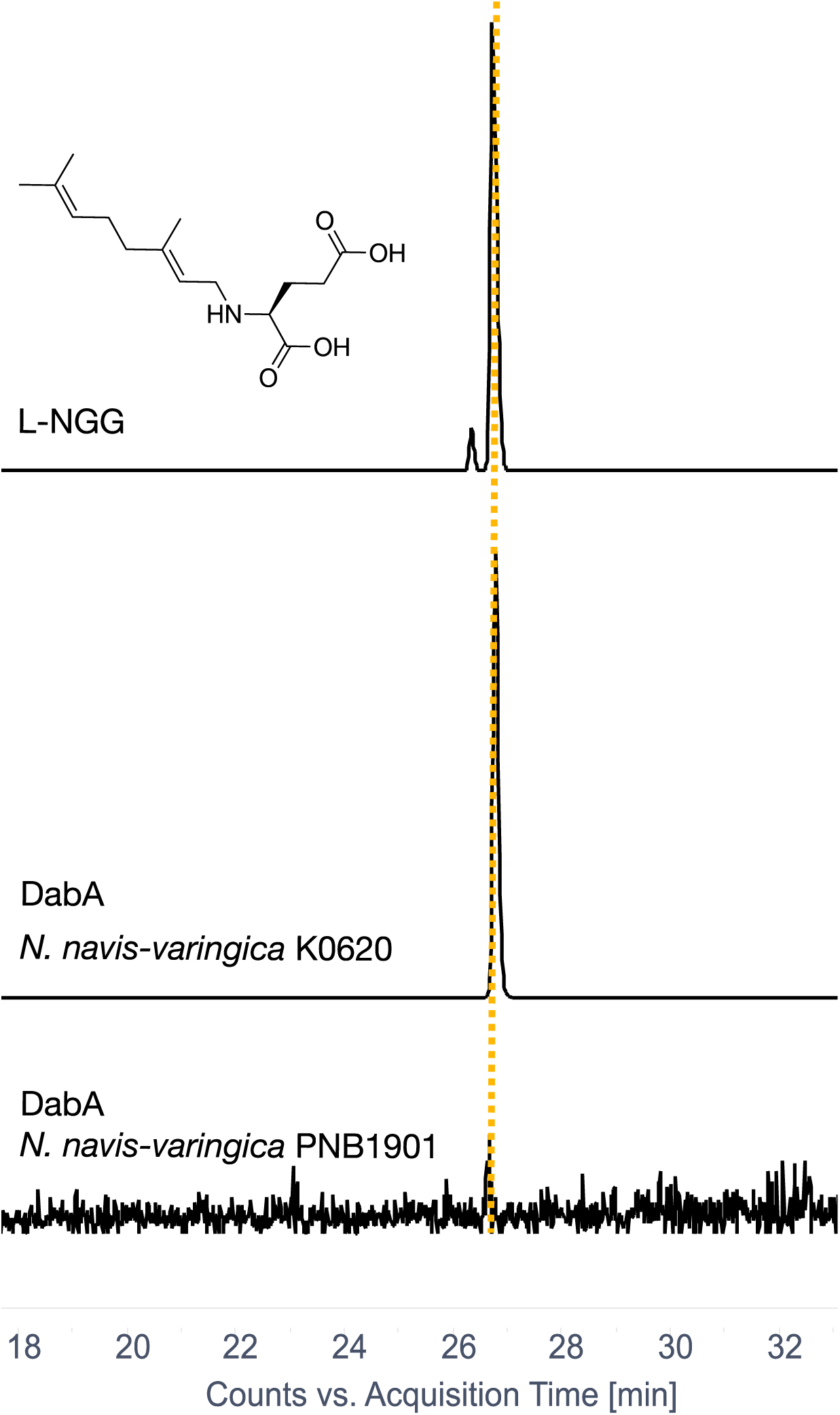
Positive ion mode LC-MS extracted ion chromatogram profiles for L-NGG (*m/z* 280.2 ± 0.1) from synthetic standard and DabA assays.

**Figure S13.**
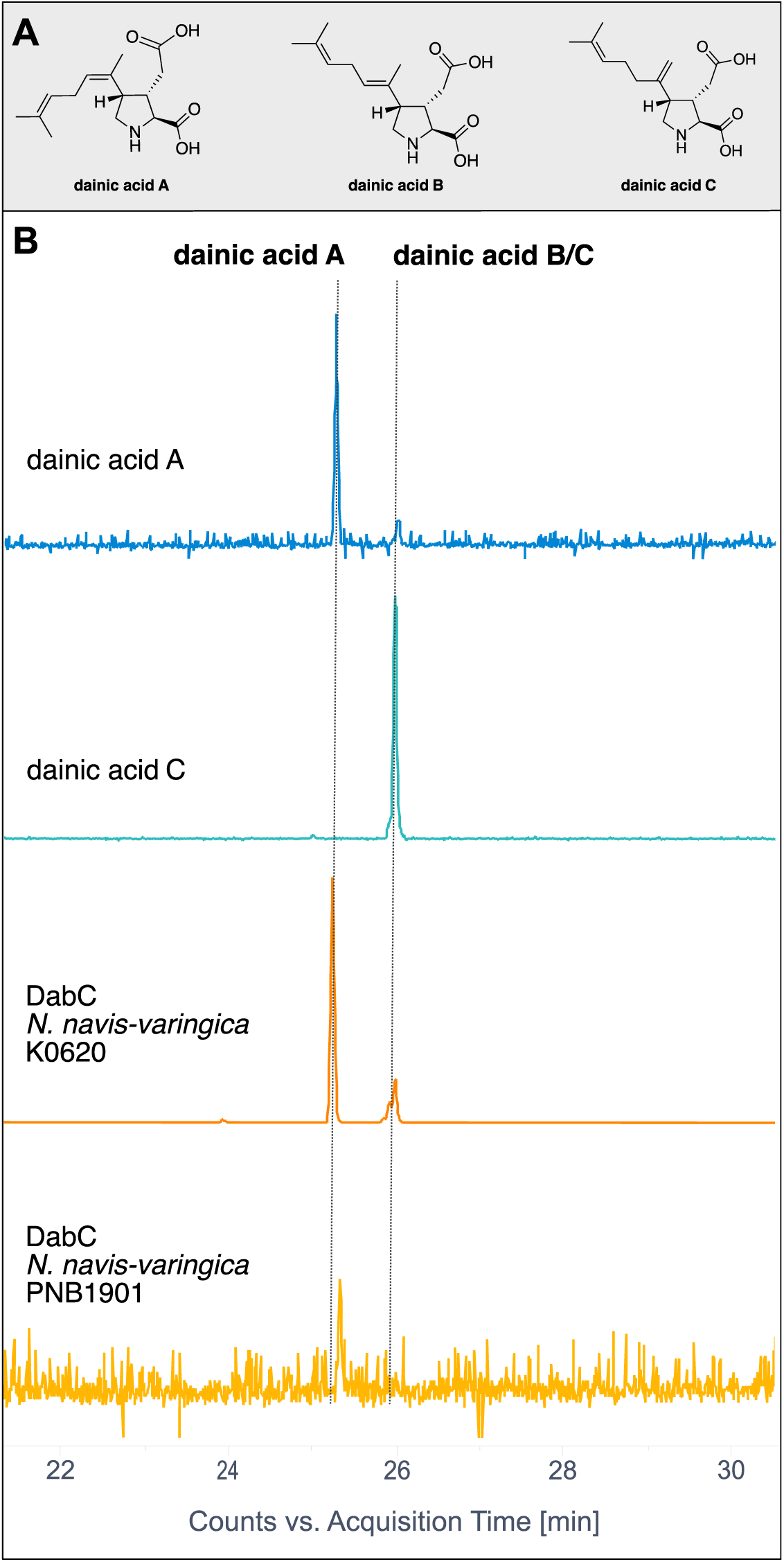
(A) Dainic acid isomers. (B) Positive ion mode LC-MS extracted ion chromatogram profiles for dainic acid isomers compared with synthetic standards (*m/z* 282.2±1.0). Dainic acids B and C have been shown to co-elute using our LCMS methods (14).

**Figure S14.**
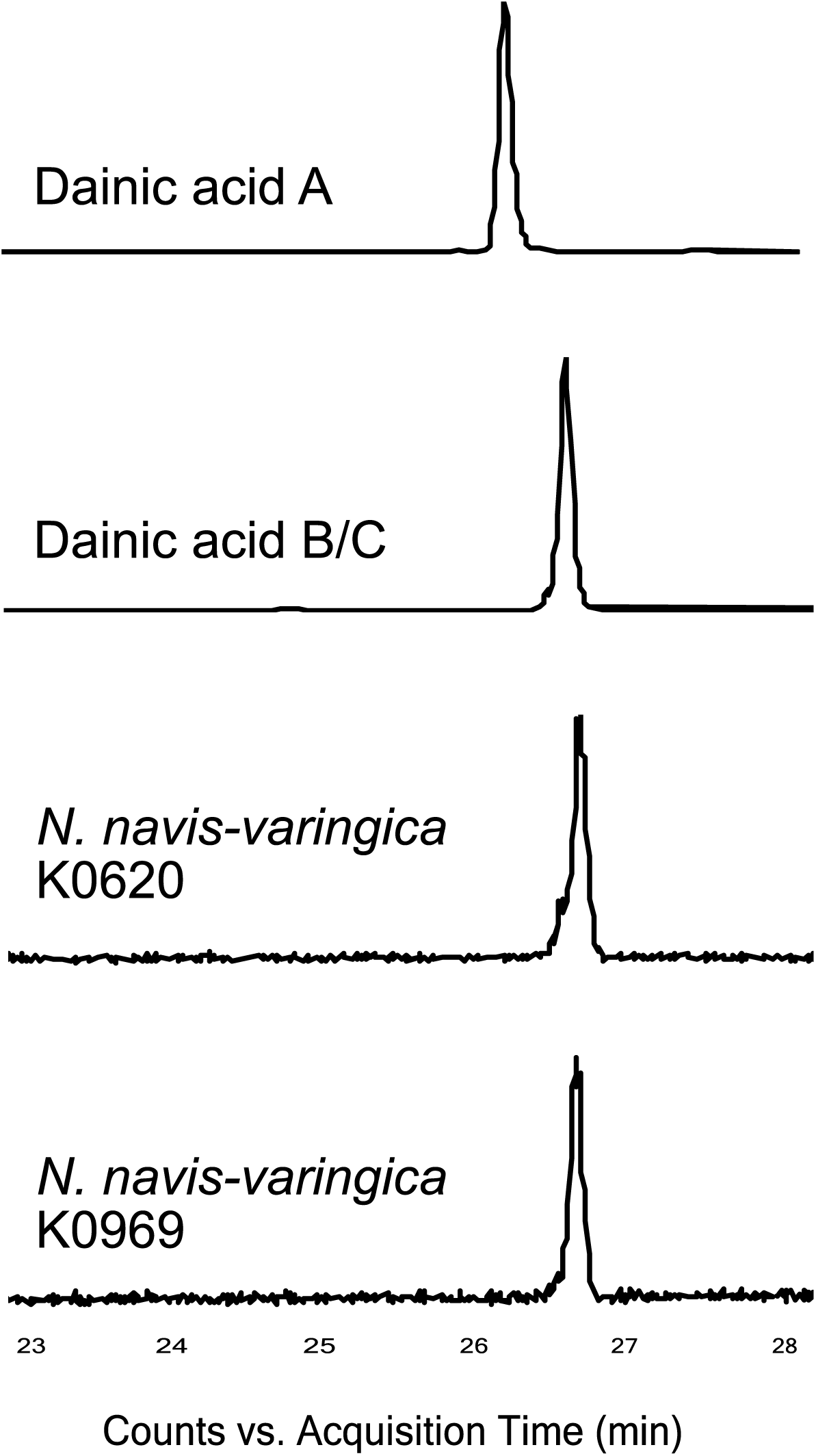
Dainic acid in culture extracts. Negative ion mode LC-MS extracted ion chromatogram profiles for dainic acid isomers compared with synthetic standards (*m/z* 280.2±0.1). Dainic acids B and C have been shown to co-elute using our LCMS methods (14).

**Figure S15.**
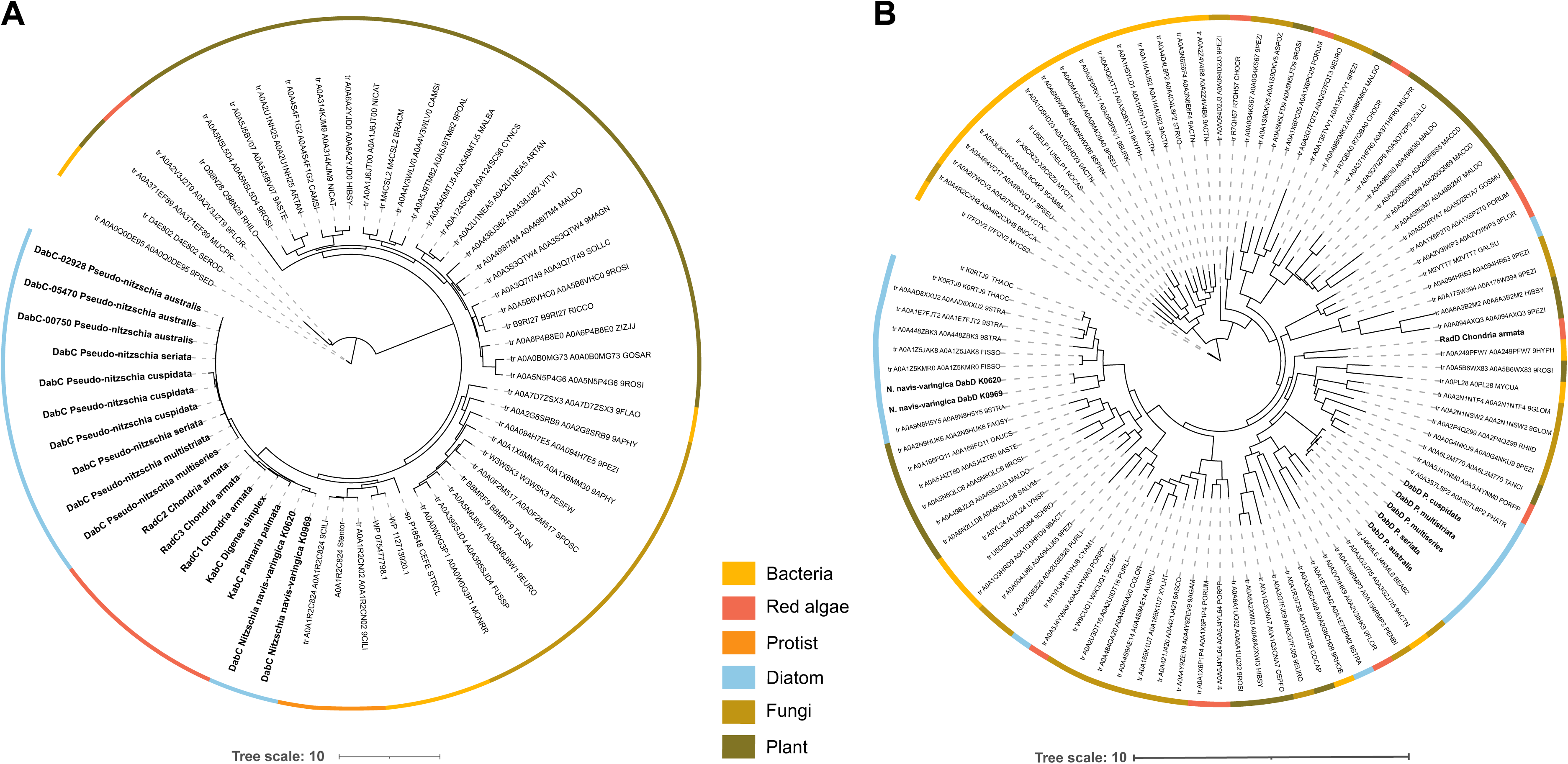
Phylogenetic analysis of (A) kainoid synthase and (B) DabD CYP450 enzymes. Maximum likelihood trees were constructed using Kalign and iTOL (EMBL). Kainoid synthase enzymes form a distinct branch, while *N. navis-varingica* DabD clusters with other diatom CYP450 sequences.

**Figure S16.**
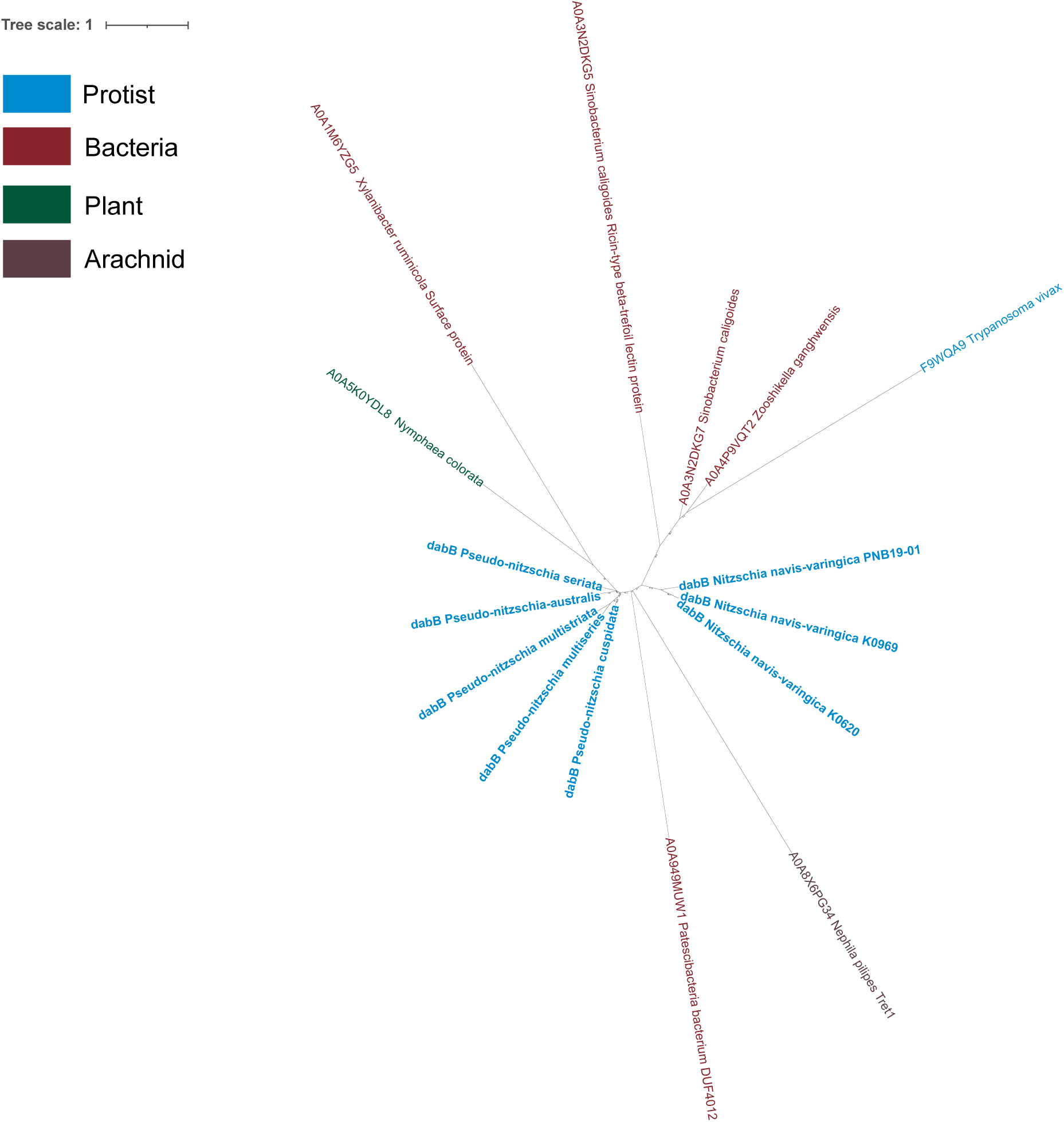
Unrooted phylogenetic analysis of DabB protein sequences.

**Figure S17.**
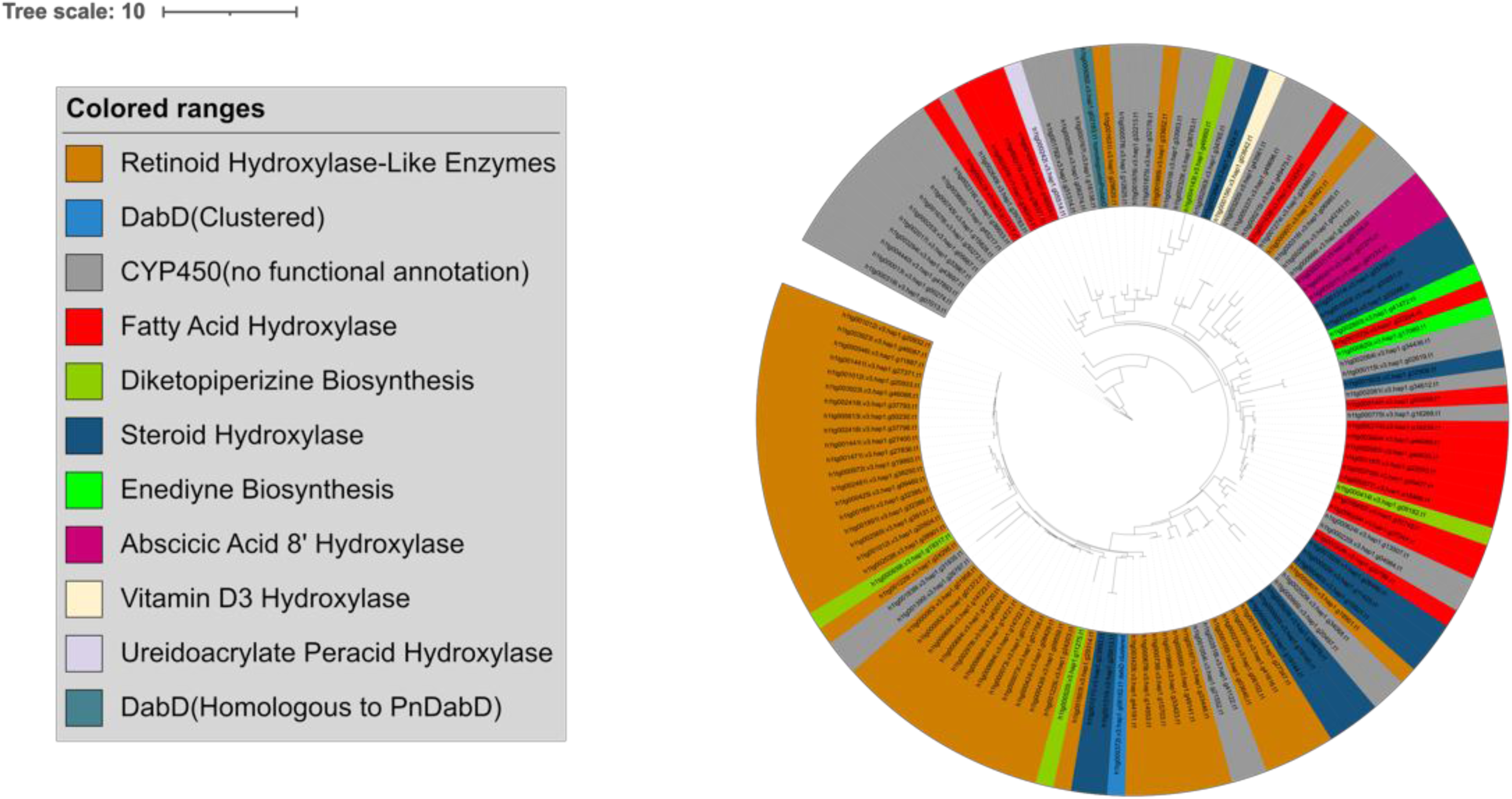
*Nitzschia navis-varingica* K0620 Cytochrome P450 phylogenetic tree. CYP function was annotated based on KEGG Orthology(KO).

**Table S3.**
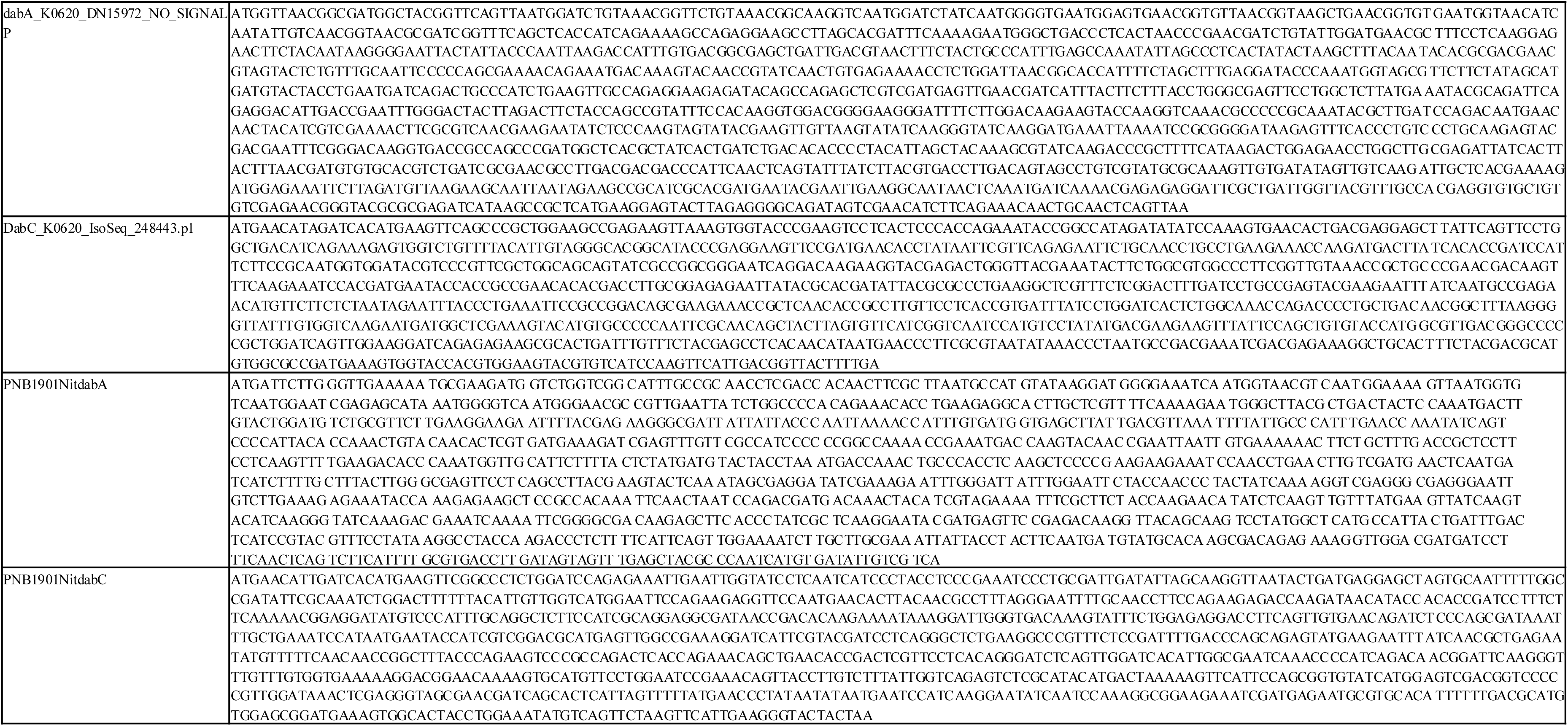
Nucleotide sequences of genes expressed *in vitro*.

